# Microbes suppress genetic disorders of hyper-activated Ras by inhibiting iron-mediated growth factor modulation in *C. elegans*

**DOI:** 10.1101/2025.09.30.679430

**Authors:** Minghui Du, Yangyang Wu, Guanqun Li, Hongyun Tang

## Abstract

Gut bacteria play crucial roles in promoting host health, but their ability to suppress the effects of genetic defects remains unclear. *Ras(gain-of-function, gf)* mutations are involved in various developmental disorders, thus identifying bacterial-activity changes capable of repressing the effects of hyper-activated Ras is important. Here, we screened mutations of all non-essential *E. coli* genes and identified 151 *E. coli* mutants that mitigate *let-60/Ras(gf)-*induced vulval developmental abnormalities in *C. elegans*. Interestingly, bacteria with mutations in genes involved in iron acquisition suppress host *Ras(gf)*-induced vulval defects through elevating 2,3-dihydroxybenzoic acid, a bacterial siderophore that sequesters iron. Consequently, host mitochondrial iron availability is decreased, which prompts the nuclear accumulation of the chromatin modifier LIN-65 and then downregulates *lin-3/EGF* transcription to repress *Ras(gf)*-induced vulval defects. Our findings identify a mechanism for coordinating Ras growth signaling with iron availability, through which gut bacteria suppress host *Ras(gf)*-induced defects and exemplify the potential of modifying gut bacterial activity to improve genetic diseases.

## Introduction

Throughout an animal’s life, the activity of gut bacteria changes, as do the composition and structure of the microbial community, all of which can impact the host’s physiology. Emerging evidence highlights the crucial roles of gut microbes in promoting animal health; the effects of these microbes range from the prevention of metabolic disorders such as diabetes to protection against infections [1–5]. The diversity of the gut microbiota represents an intriguing avenue for exploring potential therapies for various disorders. However, it remains unclear whether these microorganisms can override the effects of genetic mutations to suppress host developmental defects. Identifying microbes capable of suppressing the genetic mutation-caused developmental abnormalities holds potential for the treatment of genetic diseases. One particularly impactful type of gene mutation is RAS hyperactivation, known as *RAS gain-of-function (gf)* mutations. Germline *RAS(gf)* mutations are known to cause severe developmental disorders like Costello syndrome affecting multiple organs, while somatic *RAS(gf)* mutations are present in approximately 30% of all human cancers [6, 7]. Therefore, developing effective strategies to inhibit RAS signaling is crucial. Identifying microbes that can suppress the phenotype of *RAS(gf)* mutations could provide a new perspective for developing approaches to target RAS signaling and serve as a ‘proof of concept’ for using microbes to alleviate genetic diseases.

The RAS signaling pathway is a critical therapeutic target due to its involvement in various disorders [6, 8, 9], thus comprehensive understanding the regulation of Ras is important. In particular, it has long been speculated that RAS signaling activity may be coordinated with nutrient levels to support the role of RAS signaling in promoting cell proliferation and differentiation [6]. However, how nutrient levels are transduced into Ras signaling activity remains underexplored. For example, while iron is a crucial nutrient for cell viability, it is not yet known whether RAS signaling activity is coordinated with iron levels, and the mechanism by which the host perceives and translates iron levels to affect RAS signaling remains unclear. Gut microbes have a significant impact on host nutrient levels [10–13]; thus, exploring the potential roles of these microorganisms in shaping RAS signaling offers an exciting perspective to address the molecular basis underlying the transduction of nutrient levels into Ras signaling and their implications for animal health.

*C. elegans* has emerged as an exceptional system for studying the interactions between hosts and microbes due to its well-established genetics and the ease with which it can be rendered germ-free [14, 15]. Moreover, *C. elegans* is an excellent model for studying the regulation of Ras signaling, and previous studies on the developmental role of *let-60*/Ras in vulval precursor cell (VPC) specification have led to the discovery of highly conserved components and mechanisms of Ras signaling [16]. Specifically, the RTK-Ras-ERK pathway responds to extracellular LIN-3/EGF signals to determine whether VPCs, including six polarized epithelial cells designated P3.p to P8.p, undergo vulval or epidermal differentiation [17]. In wild-type worms, only three VPCs (P5.p, P6.p, and P7.p) are induced and form a single invagination, eventually developing into a normal vulva; however, gain-of-function (gf) mutations in *let-60/RAS*, such as the *let-60(n1046, G13E)* that mimics a Ras codon 13 mutations and often occurs in human cancers [8, 18], induce more than three VPCs, leading to the development of multiple invaginations and eventually the formation of the multiple vulva (Muv) [18, 19]. The strong conservation of LET-60/Ras signaling in *C. elegans* suggests that elucidating how bacteria regulate Ras in this organism could reveal previously unknown mechanisms of Ras signaling modulation and provide insights into the use of bacteria to suppress *RAS (gf)*-related disorders in mammals.

In this study, we aimed to uncover alterations in bacterial activities capable of repressing host hyperactivated Ras signaling. Given that inactivating bacterial genes can alter bacterial activity, the Keio library contains 3985, the all nonessential single-gene deletions in *E. coli* which represents a vast collection of bacteria with diverse types of activity changes [20]. Therefore, we employed the Keio library to identify bacterial mutants that can suppress the Muv phenotype caused by *let-60(gf)* in *C. elegans*. We identified 151 *E. coli* mutants that suppressed the Muv phenotype in *let-60(gf)* worms. These bacterial mutants included those with mutations in genes involved in various biological processes, including bacterial iron absorption. Mechanistically, microbial mutants lacking genes associated with iron absorption could enhance the production of a specific type of siderophore and then reduce the availability of mitochondrial ferrous iron in the host. The epigenetic factor Lin-65 can respond to this mitochondrial iron reduction by translocating into the nucleus, which downregulates *lin-3/EGF* expression and then inhibits Ras signaling. Therefore, our study identifies the mechanism underlying the coordination between iron levels and RAS growth signaling. Importantly, our findings provide a comprehensive understanding of suppressing the phenotype of hyperactivated Ras mutants by modulating bacterial activity, shedding light on the potential use of microbes in managing *RAS(gf)*-related disorders.

## Results

### A genome-wide screen identifies *E. coli* genes, whose inactivation suppresses the Muv phenotype in host *C. elegans* with a hyperactive *let-60/RAS* mutation

Overactivated Ras signaling results in Muv phenotype, displaying two or more ventral protrusions in each adult hermaphrodite worm [21]. To systematically investigate whether modulating bacterial activity could suppress the effects of genetic defects caused by *Ras(gf)*, we conducted a genome-wide screen using the Keio library to identify *E. coli* genes whose inactivation could suppress the Muv phenotype in *let-60(n1046, gf)* worms (Figure 1A). When administered the Keio library parental wild-type *E. coli* strain (BW25113) or OP50 *E. coli*, which is commonly used in the laboratory to feed *C. elegans*, approximately 90% of the *let-60(n1046)* animals displayed the Muv phenotype (Figure S1A), consistent with previously reported findings [18, 22]. In contrast, the wild-type N2 control worms displayed a smooth ventral surface and did not exhibit the Muv phenotype (Figure S1A). Remarkably, our screen revealed a total of 151 *E. coli* gene deletions that significantly suppressed the Muv phenotype in *let-60(n1046)* worms (Figure 1B and Table 1).

**Figure 1.**
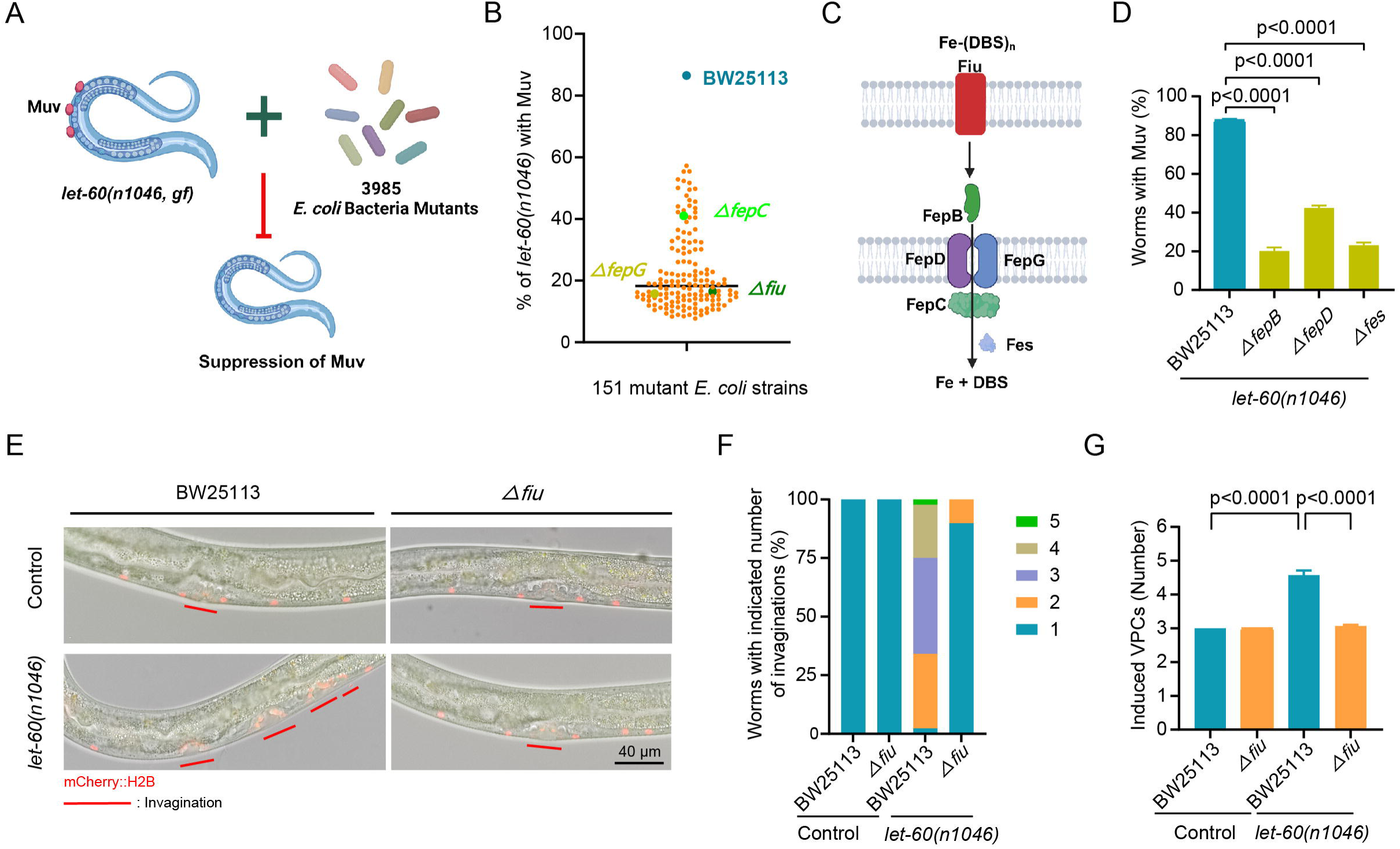
Bacterial activity changes, including the inactivation of *E. coli* iron-acquisition genes, suppressed the Muv phenotype in *let-60(n1046)* worms. **(A)** Schematic depiction of the process used to screen for bacteria that suppress the Muv phenotype in *let-60(n1046)* worms. The *E. coli* Keio Collection, consisting of 3985 single-gene knockouts, was used, and the percentage of worms with the Muv phenotype was scored on day 1 adulthood. **(B)** A total of 151 *E. coli* mutants, including those carrying mutations in the *fiu*, *fepC*, and *fepG* genes, which are involved in iron-siderophore uptake, reduced *let-60(n1046)*-induced Muv to different degrees. BW25113, the parental strain of the *E. coli* mutants, served as the control. Each dot represents the Muv rate in day-1 adult *let-60(n1046)* worms treated with the specified *E. coli* mutants, detailed in Table 1. The screening was conducted independently three times. **(C)** Diagram showing the Fe-siderophore uptake process in *E. coli* and the proteins that are involved. **(D)** Bar graph showing that bacterial mutations in other Fe-siderophore transport genes including *fepB*, *fepD* and *fes* suppress *let-60(n1046)*-induced Muv. The data are presented as the mean ± SEM. p < 0.0001 by one-way ANOVA. **(E-F)** Representative images and bar graph showing the repression of invagination formation by Δfiu bacteria treatment. (E) The *arTi85 [lin-31p::ERK::KTR::mClover::T2A::mCherry::his-11::unc-54 3’UTR + rps-27p::NeoR::unc-54 3’UTR]* reporter was used for evaluating the number of invagination, as indicated by the red line. (F) Bar graph shows the percentage of worms with the indicated number of invaginations represented using color blocking. **(G)** Bar graph showing that the *let-60(n1046)-*induced VPCs were suppressed by the Δfiu *E. coli*. The worms treated with the BW25113 parental strain were used as the control. The data are presented as the means ± SEM. p < 0.0001 was calculated by one-way ANOVA.

**Table 1.** 151 mutant *E. coli* were identified from the screen for the ability of 3,985 *E. coli* non-essential gene mutants to suppress the Muv phenotype of *let-60(n1046)* worms.

To gain insight into the functions of the bacterial genes identified in the screen, we performed Gene Ontology analyses. The *E. coli* genes, deleted in the bacterial clones capable of suppressing the Muv phenotype, were enriched in various processes, including electron transport, DNA repair, and iron homeostasis (Figure S1B and S1C). The screen results indicate that inactivating a variety of bacterial processes can suppress the Muv phenotype caused by hyperactivated Ras in the host *C. elegans*, thus providing an indication of the specific bacterial activities that can be targeted to mitigate the effects of Ras hyperactivation in the host animal.

### Inactivating the bacterial iron acquisition process renders the ability of bacteria to suppress *let-60(gf)*-induced Muv in the host

To decipher how changes in bacterial activity suppress Ras(gf)-induced Muv, we focus on understanding how inhibition of bacterial iron acquisition genes, such as *fiu*, *fepC*, and *fepG* (Figure 1B and 1C), affects host Ras signaling, given the evident enrichment of these genes in our screen. *E. coli* secretes siderophores, such as the catecholate dihydroxybenzoylserine (DBS), which form complexes with ferrous iron that are transported into the periplasm by Fiu [23]. Then, these complexes are imported into the cytoplasm through an ABC transporter [24, 25]. Once in the cytoplasm, Fes hydrolyzes ferric-DBS, releasing iron for bacterial utilization (Figure 1C). To further investigate the effect of this iron-siderophore transport process on host Ras signaling, we tested mutations in all bacterial genes in this iron acquisition process. Strikingly, we found that inactivating the other three genes involved in the Fe-DBS transport process could also significantly suppress Muv in *let-60(n1046)* worms (Figure 1D).

Given its strong Muv-suppression effect, the *E. coli* with a *fiu* gene deletion (Δ*fiu*) was used to further investigate the mechanism by which inhibiting the *E. coli* Fe-DBS transportation process suppresses *let-60(gf)*-induced Muv in the *C. elegans* host. Ras signaling determines which VPCs adopt a vulval fate at the L2-L3 stage, after which these cells undergo cell division and morphogenesis, including invagination, to form the vulva; thus, we assessed the induction of VPCs to a vulval fate and the number of invaginations to further evaluate the influence of these bacterial mutations on host Ras signaling. The *arTi85* transgene that expressed a mCherry::HIS-11 specifically in the VPCs was employed to visualize invagination [17]. We found that when *arTi85* worms were treated with wild-type BW25113 bacteria, one invagination (Figure 1E and 1F) and three induced VPCs were observed (Figure 1G), consistent with previous observations in N2 wild-type worms [26]. However, when *let-60(n1046);arTi85* worms were treated with BW25113 bacteria, more than 90% of exhibited more than one invagination (Figure 1E and 1F), and the average number of VPCs that adopted a vulval fate (induced VPCs) per worm was nearly five (Figure 1G). Strikingly, upon treatment with the Δ*fiu* mutant bacteria, the number of *let-60(n1046);arTi85* worms with more than one invagination decreased to less than 10% (Figure 1E and 1F), and the number of induced VPCs decreased to approximately three (Figure 1G). Notably, the Δfiu strain repressed only the pseudovulvas and not the normal vulva (Figure 1E-G). Together, these results indicate that the Δfiu *E. coli* can repress the *let-60(n1046)*-mediated induction of extra VPCs to a vulval fate.

Next, we investigated whether treatment with the Δ*fiu E. coli* could downregulate Ras signaling pathway activity in the *let-60(n1046)* worm host, by examining the activity of MPK-1/ERK that acts downstream of Ras [27]. MPK-1 activity in the VPCs was assessed using a previously established kinase translocation reporter, the ERK-nKTR sensor expressing *arTi85[lin-31p::ERK-KTR-mClover::T2A::mCherry-H2B]*, by calculating the nuclear Red/Green Ratio [17]. When MPK-1 kinase is inactive, the KTR remains unphosphorylated and predominantly nuclear; upon activation of the MPK-1 kinase, phosphorylation of the KTR leads to its cytoplasmic translocation, thus the nuclear Red/Green Ratio positively relates to the activity of MPK-1. The nuclear Red/Green ratio in every single VPC was scored at the L2-L3 stage when the VPCs vulva cell fate is induced. We found that Δ*fiu E. coli* treatment is able to suppress the ERK activity in the VPCs of *let-60(n1046)* worms, compared to the BW25113 treatment (Figure S1D). These results support that the Δ*fiu* bacteria treatment decreases Ras signaling pathway activity to block the extra vulva fate induction and to suppress Muv. Moreover, we observed that Δfiu *E. coli* also suppressed the Muv phenotype induced by two other Ras gain-of-function alleles, *let-60(n1700)* (Figure S1E) and *let-60(ga89)* (Figure S1F). These results together indicate that the Muv-suppressive effect of the Δfiu *E. coli* strain is not allele specific, thus further supporting the contention that the Δfiu *E. coli* strain inhibits the Ras signaling pathway. In summary, inactivating bacterial genes involved in siderophore-iron import suppresses the developmental abnormalities caused by hyperactivated Ras signaling.

### **Δ***fiu E. coli* promotes 2,3-DHBA siderophore production to reduce host iron availability, thereby suppressing Muv induced by the hyperactivated Ras

Next, we aimed to investigate the mechanism through which the Δfiu *E. coli* strain suppressed the *let-60(n1046)*-induced Muv phenotype. As the siderophore transportation is the main process for bacteria iron acquisition [25], we postulated that inactivating the Fe-(DBS)_n_ transport process in bacteria may boost the production of siderophore for reestablishing bacteria’s ability to acquire iron, which may thus lead to reduced Ras signaling in the host. Indeed, we found that compared with BW25113, Δ*fiu* bacteria secreted more 2,3-DHBA, a DBS siderophore, as evidenced by its higher abundance in the Δ*fiu* bacteria culture medium (Figure 2A). Importantly, we revealed that supplementation with 2,3-DHBA is able to dramatically repress *Ras(gf)-*induced Muv, evidenced by the Muv phenotype in *let-60(n1046)* worms grown on BW25133 strain reduced from approximately 90% to 10% (Figure 2B). These results support the role of increased production of 2,3-DHBA by Δ*fiu* bacteria in suppressing *Ras(gf)*-induced Muv.

**Figure 2.**
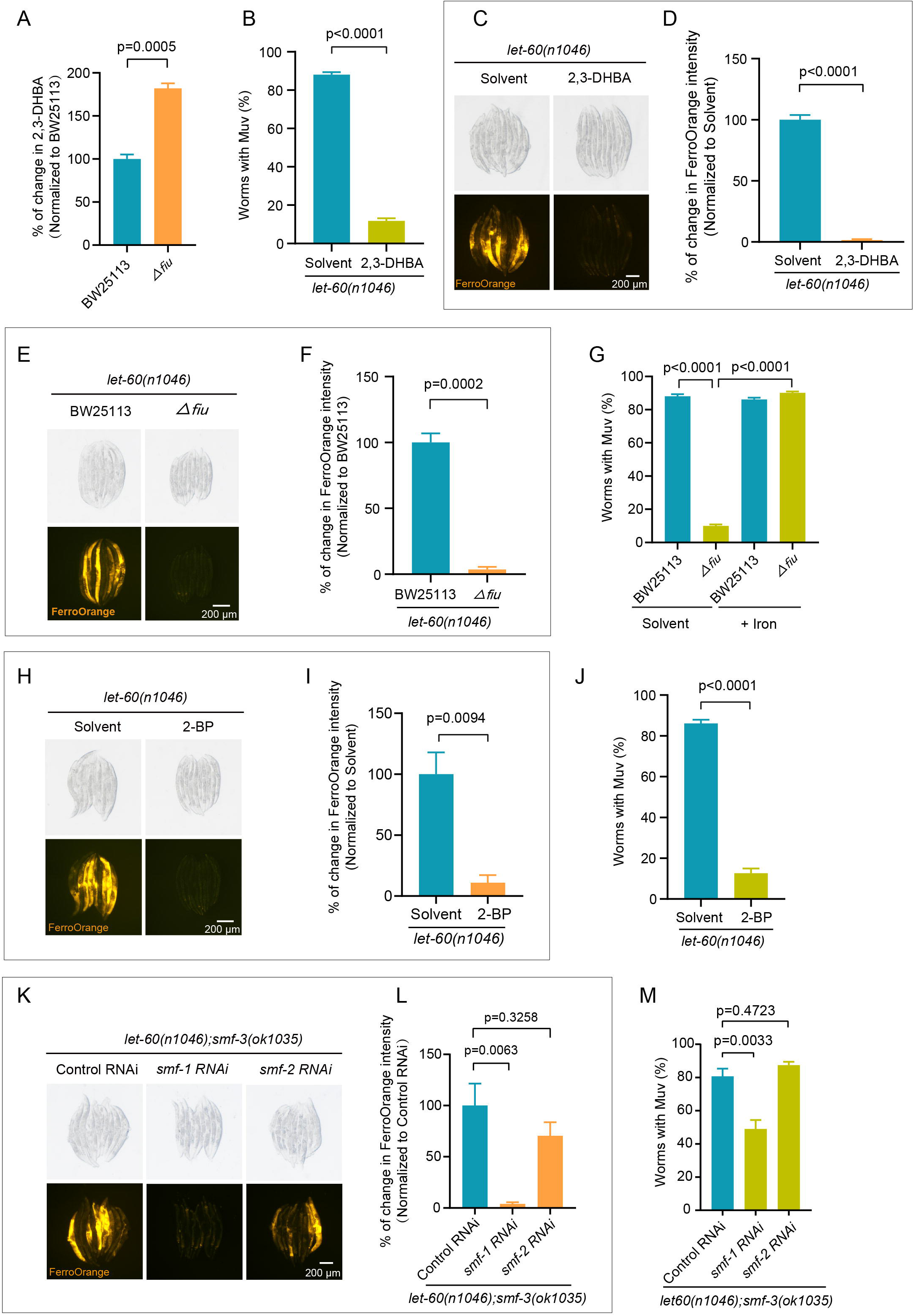
Δ*fiu E. coli* generates increased 2,3-DHBA siderophore to reduce iron availability, thereby suppressing *let-60(n1046)*-induced Muv. **(A)** A bar graph showing the percentage of 2,3-DHBA secreted by Δ*fiu E. coli* relative to that produced by BW25113. The 2,3-DHBA in the bacterial supernatant was quantified using mass spectrometry. The data are presented as the mean ± SEM by unpaired t test. **(B)** A bar graph illustrating the potent suppressive effect of 2,3-DHBA on the Muv rate in *let-60(n1046)* mutants. Worms were treated with either 18 mg/mL 2,3-DHBA or solvent, and Muv phenotypes were assessed at the adult day 1 stage. Data are presented as the mean ± SEM, determined by an unpaired t-test. **(C-D)** Representative images and a bar graph illustrate the decrease in ferrous iron, stained with FerroOrange, in *let-60(n1046)* worms treated with 2,3-DHBA. Worms treated with DMSO or 18 mg/mL 2,3-DHBA were assessed at the 1-day adult stage. Data are presented as the mean ± SEM, analyzed by unpaired t-test. **(E-F)** Images and bar graphs illustrating the decrease of ferrous iron, stained with FerroOrange, in the *let-60(n1046)* worms treated with Δ*fiu E. coli*. Intensity of FerroOrange was analyzed in the worms treated with BW25113 or Δ*fiu* bacteria at the day-1 adult stage. The data are presented as the means ± SEM by unpaired t test. **(G)** Bar graphs illustrating that iron supplementation reverses the suppressive effect of Δ*fiu* bacteria on *let-60(n1046)*-induced Muv. The Muv rate in *let-60(n1046)* worms treated with Δ*fiu* or control BW25113 strains, with or without 4 mM FeCl_3_ supplementation, was analyzed. Data are presented as means ± SEM and analyzed by one-way ANOVA. **(H-I)** Representative images and corresponding bar graphs illustrate the reduction in ferrous iron, stained with FerroOrange, in worms treated with the iron chelator reagent 2-BP. The intensity of FerroOrgane in *let-60(n1046)* worms, treated with either 2-BP or solvent, were analyzed at day-1 adult stage. Data are presented as the mean ± SEM by unpaired t test. **(J)** A bar graph showing the decrease in Muv rate in *let-60(n1046)* worms treated with 2-BP. The Muv rate of *let-60(n1046)* worms, treated with either 2-BP or solvent, was analyzed at the day-1 adult stage. Data are presented as the mean ± SEM by unpaired t test. **(K-L)** Images and bar graphs showing that ferrous iron, stained with FerroOrange, is decreased in the *let-60(n1046)* worms under *smf-3(lf);smf-1(RNAi)*. The intensity of FerroOrange in *smf-3(lf);let-60(n1046)* worms treated with *smf-1*(RNAi), *smf-2(RNAi)* or pl4440 RNAi control was analyzed at the day-1 adult stage. The data are presented as the mean ± SEM by one-way ANOVA. **(M)** Bar graphs showing that *smf-3(lf);smf-1(RNAi)* partially suppresses the *let-60(n1046)-* induced Muv rate. The Muv rate in *smf-3(lf);let-60(n1046)* worms treated with *smf-1*(RNAi), *smf-2(RNAi)* or pl4440 RNAi control was analyzed at the day-1 adult stage. The data are presented as the mean ± SEM by one-way ANOVA.

Then, we tested whether increased generation of 2,3-DHBA from Δ*fiu* bacteria may reduce host iron availability to downregulate Ras signaling, given that siderophore sequesters iron [28]. Firstly, we evaluated the labile ferrous iron levels using the dye FerroOrange, which reacts with free ferrous iron to produce a fluorescent signal [29], and found that the intracellular iron availability in *let-60(n1046)* worms treated with the BW25113 bacteria supplemented with 2,3-DHBA (Figure 2C and 2D) or treated with Δfiu mutant *E. coli* (Figure 2E and 2F) were obviously decreased, as indicated by the reduced FerroOrange fluorescence intensity following treatment with 2,3-DHBA or Δfiu mutant *E. coli*. Subsequently, we tested the potential reversal of Δfiu-mediated Muv suppression by restoring iron levels in the *let-60(n1046)* worms. During iron absorption, ferric ions undergo reduction and are imported for utilization, so we used FeCl_3_ for iron supplementation [30]. After FeCl_3_ supplementation, the Muv percentage increased from approximately 10% to approximately 90%, indicating that restoring iron completely reverses the Δfiu *E. coli*-mediated Muv suppression (Figure 2G). Therefore, Δfiu *E. coli* suppresses *let-60(n1046)*-induced Muv through downregulating host iron availability.

Furthermore, we found that treatment with iron-chelator 2,2-bipyridyl (2-BP), that sequesters iron in the *let-60(n1046)* worms (Figure 2H and 2I), can also suppress *let-60(n1046)*-induced Muv. The percentage of *let-60 (n1046)* worms with Muv decreased from 90% to 12% following 2-BP-mediated iron chelation (Figure 2J). This result further demonstrates that the suppressive effect of Δfiu *E. coli* on *let-60(gf)*-induced Muv is mediated by a reduction in labile iron. Notably, 2-BP treatment decreased FerroOrange fluorescence intensity (Figure 2H and 2I), confirming that FerroOrange can indicates free ferrous iron levels in worms. Moreover, FeCl_3_ supplementation also reverses the Muv suppression mediated by treatment with other bacterial mutants lacking Fe-(DBS)_n_ transportation-related genes (Figure S2A). Taken together, these results demonstrate that inactivation of bacterial Fe-(DBS)_n_ transport decreases available iron levels in the host, which leads to the suppression of *Ras(gf)*-induced Muv.

To confirm the role of host iron insufficiency in the inhibition of Ras(gf)-induced Muv, we experimentally decreased the ability of the worms to absorb iron and then assessed any potential suppression of Muv in *let-60(gf)* worms. *smf-1/-2/-3* encode iron transport proteins responsible for iron absorption in *C. elegans* [30]. *smf-3* deletion (*smf-3(lf)*) (Figure S2B) or RNAi against the *smf-1* or *smf-2* genes (Figure S2C) did not reduce iron absorption, as indicated by unchanged FerroOrange fluorescence intensity in the worms; accordingly, *let-60(gf)*-induced Muv was not suppressed by these treatments (Figure S2D and S2E), which suggests functional redundancy in iron transport among these genes. We next performed RNAi against *smf-1* or *smf-2* in the *smf-3(lf)* background to overcome this redundancy. Strikingly, we found that *smf-1* RNAi, but not *smf-2* RNAi, decreased labile iron levels in *smf-3(lf);let-60(n1046)* worms (Figure 2K and 2L). Consistent with the role of decreased labile iron in suppressing *let-60(n1046)*-induced Muv, partial suppression of Muv was observed in *smf-3(lf);let-60(n1046)* worms treated with *smf-1* RNAi but not in those treated with *smf-2* RNAi (Figure 2M). Taken together, the findings indicate that Δ*fiu* bacteria produces increased levels of 2,3-DHBA that reduce iron availability in *C. elegans,* thereby suppressing the hyperactivated Ras-induced Muv phenotype.

Moreover, to assess whether this effect was restricted to *E. coli*, we examined whether *Bacillus subtilis*, another naturally existed gut bacterium capable of scavenging inorganic iron from the environment [31], could also suppress the *let-60(gf)*-induced Muv phenotype through decreasing the available iron levels. Compared to the OP50 *E. coli* control, treatment with *B. subtilis* markedly suppressed Muv in adult *let-60(gf)* worms (Figure S2F). Importantly, this suppression was completely reversed by iron supplementation (Figure S2F). Thus, both *E. coli* gene mutant strains and *B. subtilis* can suppress the effects of hyperactivated Ras by decreasing the amount of labile iron in the host.

### Decreased labile ferrous iron in the mitochondria accounts for the **Δ***fiu E. coli*-mediated suppression of *let-60(gf)*-induced Muv

Given that mitochondria are major organelles that consume iron for various vital biological processes [32], we investigated whether iron decline in the mitochondria may be responsible for the suppression of *let-60(n1046)*-induced Muv by Δ*fiu E. coli*. Thus, we investigated whether Δ*fiu E. coli* reduces available iron levels in host mitochondria using Mito-FerroGreen, which specifically detects labile ferrous iron in mitochondria [33]. First, we validated the specificity of Mito-FerroGreen for labeling mitochondrial ferrous ions in worms by confirming the colocalization of Mito-FerroGreen staining with MitoTracker Deep Red, a mitochondria-specific dye (Figure S3A). Furthermore, we observed that the Mito-FerroGreen fluorescence intensity was lower in *let-60(n1046)* worms treated with Δfiu than in those treated with BW25113 and that this effect was reversed through iron supplementation (Figure 3A and 3B). This result, combined with our data showing that iron supplementation can counteract the ability of Δfiu to suppress Muv in *let-60(gf)* worms (Figure 2G), supports the notion that mitochondrial iron decrease is involved in the Δfiu-mediated suppression of *let-60(n1046)*-induced Muv. Moreover, we also observed a reduction in mitochondrial iron levels upon supplementation with the iron chelator 2-BP (Figure 3C and 3D), consistent with the role of 2-BP in suppressing *let-60(n1046)*-induced Muv (Figure 2J). Taken together, these results indicate that the presence of Δfiu *E. coli* leads to a reduction in the amount of available iron in mitochondria in *C. elegans*.

**Figure 3.**
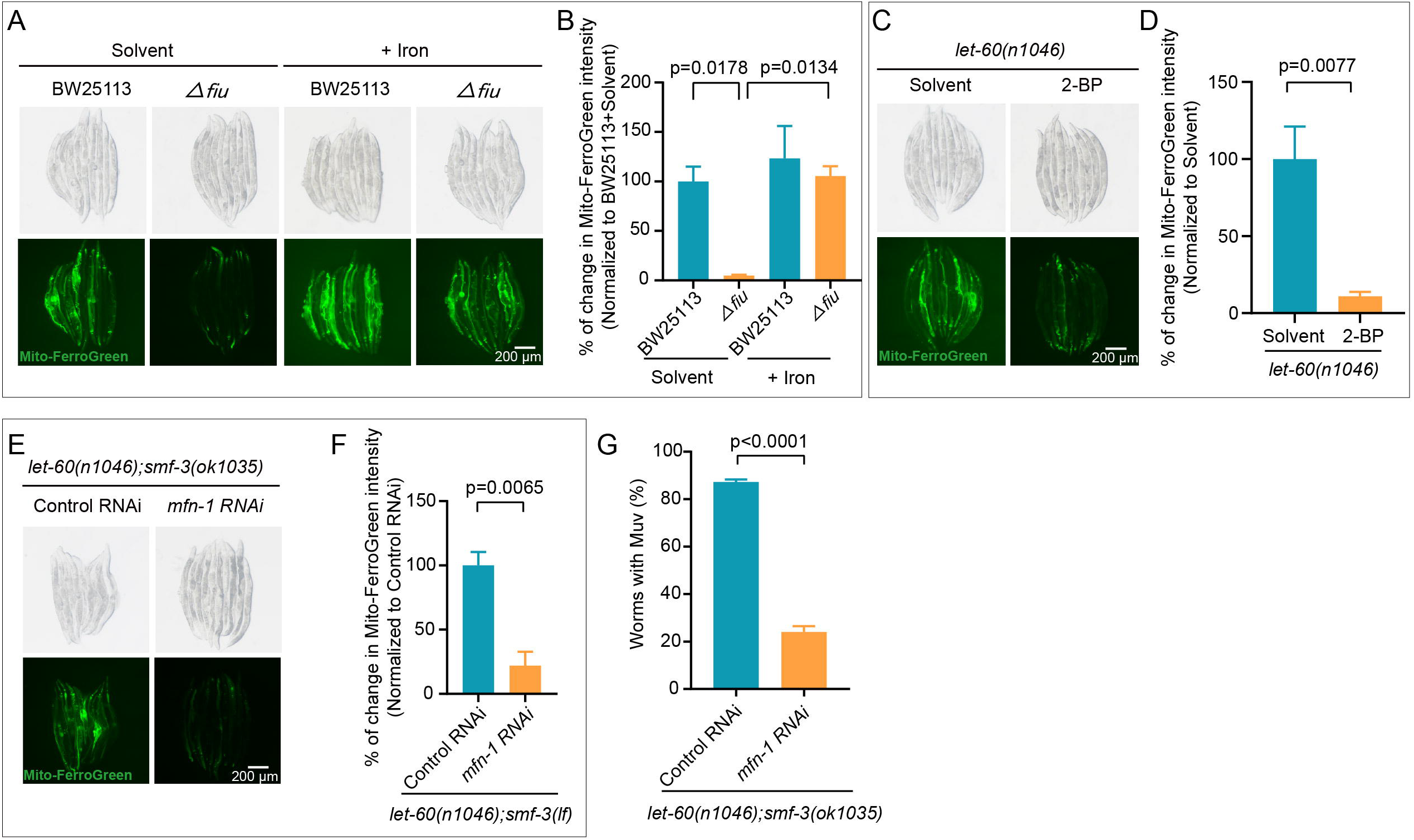
Δ*fiu E. coli* causes decreased mitochondrial iron availability, which suppresses Muv in *let-60(n1046)* worms. (**A-B**) Images and bar graphs demonstrate that mitochondrial ferrous iron, stained with Mito-FerroGreen, is reduced in *let-60(n1046)* worms treated with Δ*fiu* mutant bacteria, which is reversed by iron supplementation. The fluorescence intensity of Mito-FerroGreen in worms treated with the specified bacteria, with or without FeCl_3_ supplementation, was analyzed at the adult day-1 stage. The data are presented as the mean ± SEM by one-way ANOVA. **(C-D)** Images and bar graphs indicating that 2-BP treatment causes the decrease in mitochondrial ferrous iron, stained with Mito-FerroGreen, in *let-60(n1046)* worms. The intensity of Mito-FerroGreen is evaluated in the *let-60(n1046)* worms with 2-BP treatment at the day-1 adult stage. The data are presented as the mean ± SEM by unpaired t test. **(E-F)** Images and bar graphs showing that the levels of mitochondrial ferrous iron, stained with Mito-FerroGreen, are decreased in *let-60(n1046);smf-3(ok1035)* with *mfn-1(RNAi)* treatment. The fluorescence intensity of Mito-FerroGreen is evaluated in the *let-60(n1046);smf-3(ok1035)* worms with indicated RNAi treatment at the day-1 adult stage. The data are presented as the mean ± SEM by unpaired t test. **(G)** Bar graphs showing that the Muv rate is decreased in *let-60(n1046);smf-3(ok1035)* after *mfn-1(RNAi)* treatment. The Muv rate was evaluated in the *let-60(n1046);smf-3(ok1035)* worms with indicated RNAi treatment at the day-1 adult stage. The data are presented as the mean ± SEM by unpaired t test.

Next, we inhibited iron import into mitochondria to test the potential role of mitochondrial iron insufficiency in suppressing *Ras(gf)*-induced Muv. MFN-1 was previously implicated in mitochondrial iron import [34]; however, we observed no reduction labile iron levels in the mitochondria of worms following *mfn-1*(RNAi) treatment (Figure S3B and S3C). This result is consistent with the observation that *let-60(gf)*-induced Muv was not suppressed following *mfn-1(RNAi)* treatment (Figure S3D). Then, we attempted to decrease mitochondrial iron levels by performing *mfn-1*(RNAi) in the *smf-3*(ok1035) background, which slightly compromises worms’ ability to absorb extracellular iron. Interestingly, compared with the *mfn-1(RNAi)* control worms and *smf-3(ok1035)* control animals, the *mfn-1(RNAi);smf-3(ok1035)* group exhibited a noticeable reduction in labile iron within the mitochondria (Figure 3E and 3F, Figure S3B and S3C). Strikingly, compared to the control, *mfn-1(RNAi);smf-3(ok1035)* strongly repressed the *let-60(n1046)*-induced Muv phenotype, with the Muv rate decreasing in these worms from 90% to approximately 24% (Figure 3G). Taken together, these findings indicate that the presence of the Δfiu *E. coli* leads to a reduction in the available iron in mitochondria, which is responsible for the suppression of *let-60(gf)-* induced Muv.

### LIN-65, a mitochondrial stress response mediator, responds to **Δ***fiu E. coli-*induced decrease in labile ferrous iron, which partially suppresses *let-60(gf)*-induced Muv

Given that Δ*fiu E. coli* affects host mitochondrial iron homeostasis, which is crucial for maintaining normal mitochondrial function, we next tested whether a mitochondrial stress response may be triggered by Δ*fiu E. coli*, potentially leading to the suppression of *let-60(gf)-* induced Muv. We first examined whether a host mitochondrial stress response was induced in *let-60(n1046)* worms treated with Δfiu *E. coli* by evaluating the expression of *hsp-6p::GFP*, which has been previously shown to be activated during mitochondrial stress [35], and assessing its responsiveness to changes in iron levels. Indeed, treatment with Δfiu *E. coli* induced *hsp-6p::GFP* expression, and this activation could be reversed by iron supplementation (Figure S4A and S4B). Additionally, treatment with the iron chelator 2-BP also mimicked the impact of Δfiu *E. coli* on activating *hsp-6p::GFP* expression (Figure S4C and S4D). These findings indicated that the Δfiu-induced reduction in available iron activated the mitochondrial stress response.

Next, we explored whether activation of the mitochondria response mediated the suppression of the *let-60(gf)*-induced Muv phenotype by the Δ*fiu* strain. Initially, we investigated the potential involvement of ATFS-1, a master regulator required for activating the mitochondrial unfolded protein response (UPR^mito^) [35]. We found that an *atfs-1(et15, gf)* mutation, that activated the UPR^mito^ [36], was unable to suppress the *let-60(n1046)*-induced Muv phenotype (Figure S4E). Conversely, an *atfs-1(gk3094, lf)* mutation, that repressed the UPR^mito^ [37], did not reverse the suppressive effect of the Δ*fiu E. coli* on the *let-60(n1046)*-induced Muv (Figure S4F). Thus, ATFS-1 unlikely involves in the suppression of *let-60(n1046)*-induced Muv phenotype by Δ *fiu E. coli*. Then, we investigated the potential involvement of *lin-65*, which encodes a nuclear cofactor and responds to mitochondrial stress independent of *atfs-1* [38]. We found that the percentage of Δfiu *E. coli*-treated *let-60(n1046)* worms with Muv increased from approximately 12% to 60% in the presence of the *lin-65(lf)* mutation (Figure 4A), indicating that loss of *lin-65* function partially reversed the suppressive effect of Δfiu bacteria on the *let-60(n1046)*-induced Muv phenotype. Similarly, the suppressive effect of 2-BP on *let-60(n1046)*-induced Muv was largely reversed by *lin-65(lf)* (Figure 4A). These results support that LIN-65 activation mediates the suppressive impact of Δfiu bacteria and iron decrease on *let-60(gf)*-induced Muv.

**Figure 4.**
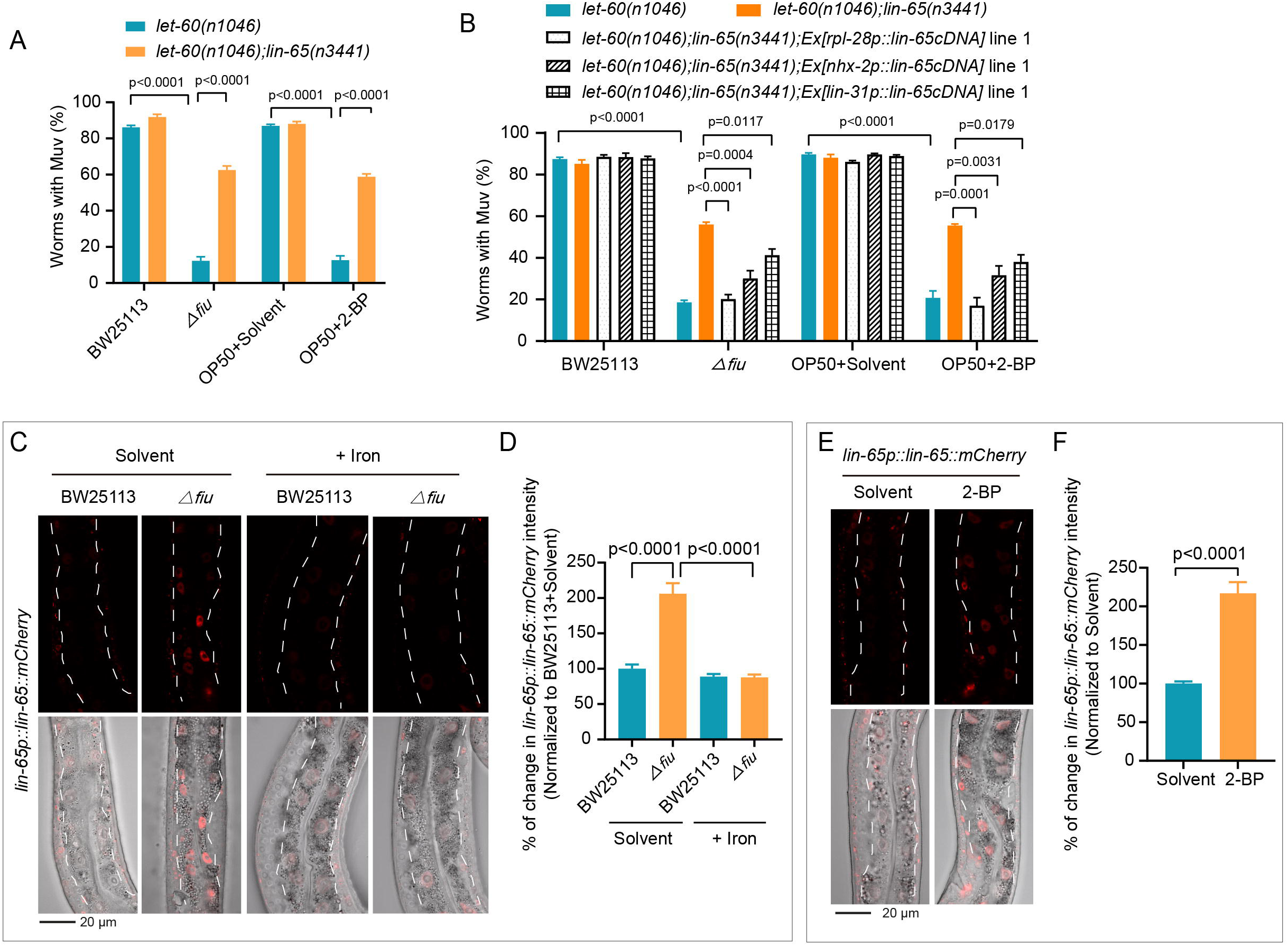
LIN-65 exhibits increased nuclear localization in response to bacteria-induced iron deficiency, partially mediating the suppressive effect of Δ*fiu E. coli* on the Muv phenotype of *let-60(n1046)* worms. **(A)** Bar graphs illustrating that the suppressive effect of Δ*fiu* bacteria and 2-BP on *let-60(n1046)*-induced Muv partially relies on *lin-65*. The Muv rate in *let-60(n1046)* and *let-60(n1046);lin-65(n3441)* worms, with specified treatments, was analyzed at the day-1 adult stage. The data are presented as the mean ± SEM by one-way ANOVA. **(B)** Bar graphs indicate the Muv rate of *let-60(n1046);lin-65(n3441)* worms carrying the specified extra-chromosomal array transgenes and treated with the designated bacteria and 2-BP. The promoters *rpl-28p*, *nhx-2p*, and *lin-31p* were used to express *lin-65* cDNA in ubiquitous tissues, the intestine, and VPCs, respectively. The data are presented as the mean ± SEM by one-way ANOVA. **(C-D)** Representative images and bar graphs demonstrate the increase in LIN-65::mCherry fluorescence in the intestinal nuclei of worms treated with Δ*fiu E. coli*, an effect reversed by iron supplementation. The mCherry signal was analyzed in *let-60(n1046);Is[lin-65p::LIN-65::mCherry]* worms exposed to the indicated bacteria, with or without FeCl_3_ supplementation. The percentage is determined by normalized to the worms treated with both BW25113 control bacteria and solvent. The dotted line delineating the intestine. The data are presented as the mean ± SEM by one-way ANOVA. **(E-F)** Representative images and bar graphs showing that iron chelator 2-BP treatment increases the nuclear localization of LIN-65::mCherry. The mCherry signal was analyzed in *let-60(n1046);Is[lin-65p::LIN-65::mCherry]* worms subjected to the indicated treatments. The dotted line depicts the intestine. The data are presented as the mean ± SEM by one-way ANOVA.

To determine the tissue in which LIN-65 expression mediates Muv suppression, transgenes expressing *lin-65* in different tissues were constructed. When LIN-65 was expressed via the ubiquitous *rpl-28* promoter in *let-60(n1046);lin-65(lf)* worms, the percentage of worms with the Muv phenotype decreased from approximately 56% to below 20% upon Δfiu and 2-BP treatment (Figure 4B and S4G). This decrease was almost equal to that observed in *let-60(gf)* worms treated with Δfiu or 2-BP (Figure 4B and S4G). This result further demonstrated that the effects of Δfiu bacteria and iron insufficiency in suppressing *let-60(gf)*-induced Muv are mediated by LIN-65. We then constructed transgenes to specifically express LIN-65 in the intestine and VPCs in *let-60(gf);lin-65(lf)* worms via the *nhx-2p* and *lin-31p* promoters, respectively. Transgenes expressing *lin-65* in either the intestine or VPCs partially inhibited the ability of *lin-65(lf)* to reverse the Δfiu and 2-BP-mediated suppression of *let-60(gf)*-induced Muv (Figure 4B and S4G). The stronger reversal effect observed following ubiquitous *lin-65* expression than observed following tissue-specific expression suggests that *lin-65* functions in multiple tissues to suppress Ras signaling in VPCs.

We further investigated whether LIN-65 responds to the presence of Δfiu *E. coli* and to 2-BP-mediated iron reduction. To detect the subcellular localization of LIN-65, a previously established transgene, *Is[lin-65p::LIN-65::mCherry]* [38], was used. We found that LIN-65 was expressed in the intestine (Figure 4C and 4E). Interestingly, compared to treatment with the control bacteria BW25113 or the solvent control, treatment with Δfiu bacteria or 2-BP increased LIN-65::mCherry levels in the intestinal nuclei of *let-60 (n1046)* worms (Figure 4C-4F). Importantly, this increase of LIN-65 in the intestinal nuclei was reversed by iron supplementation (Figure 4C-4D). Thus, LIN-65 responds to reductions in available iron, including those induced by bacteria. Notably, LIN-65 was also known as a synMuv B gene required for normal vulval development [39], further supporting the role of LIN-65 activation in suppressing Ras signaling. In summary, our findings indicate that the LIN-65 crucially mediates the role of Δfiu bacteria-mediated decrease in available iron levels in the suppression of *Ras(gf)*-induced Muv.

### In response to presence of Δ*fiu E. coli*, LIN-65 represses *lin-3/EGF* transcription to decrease *let-60(n1046)*-induced Muv

Next, we aimed to understand how *lin-65* represses *Ras(gf)*-induced Muv. Interestingly, previous studies have found that the upstream EGF receptor is necessary for certain *Ras(gf)* mutations to activate downstream signaling cascades [40, 41]. Our data, shown above, indicate that intestinal LIN-65 can counteract Ras signaling in VPCs (Figure 4B and S4G), suggesting the potential involvement of a cell non-autonomous regulation by the ligand LIN-3/EGF. Therefore, we postulated that a decrease in LIN-3 may mediate LIN-65-mediated suppression of Ras(gf) signaling in VPCs. To test this, we examined the potential of *lin-3* repression in suppressing Muv induced by downstream hyperactivated Ras by performing *lin-3 (RNAi)* in *let-60(gf)* worms, as loss-of-function mutations in *lin-3* cause severe defects. Indeed, downregulating *lin-3* by RNAi knockdown resulted in partial suppression of the *let-60(n1046)*-induced Muv phenotype, as indicated by a decrease in the Muv percentage from 90% to 68% (Figure 5A). This ligand-dependent effect of Ras(gf) aligns with previous studies showing that the activity of certain *Ras(gf)* mutations is at least partially dependent on the upstream EGFR activity [40, 41].

**Figure 5.**
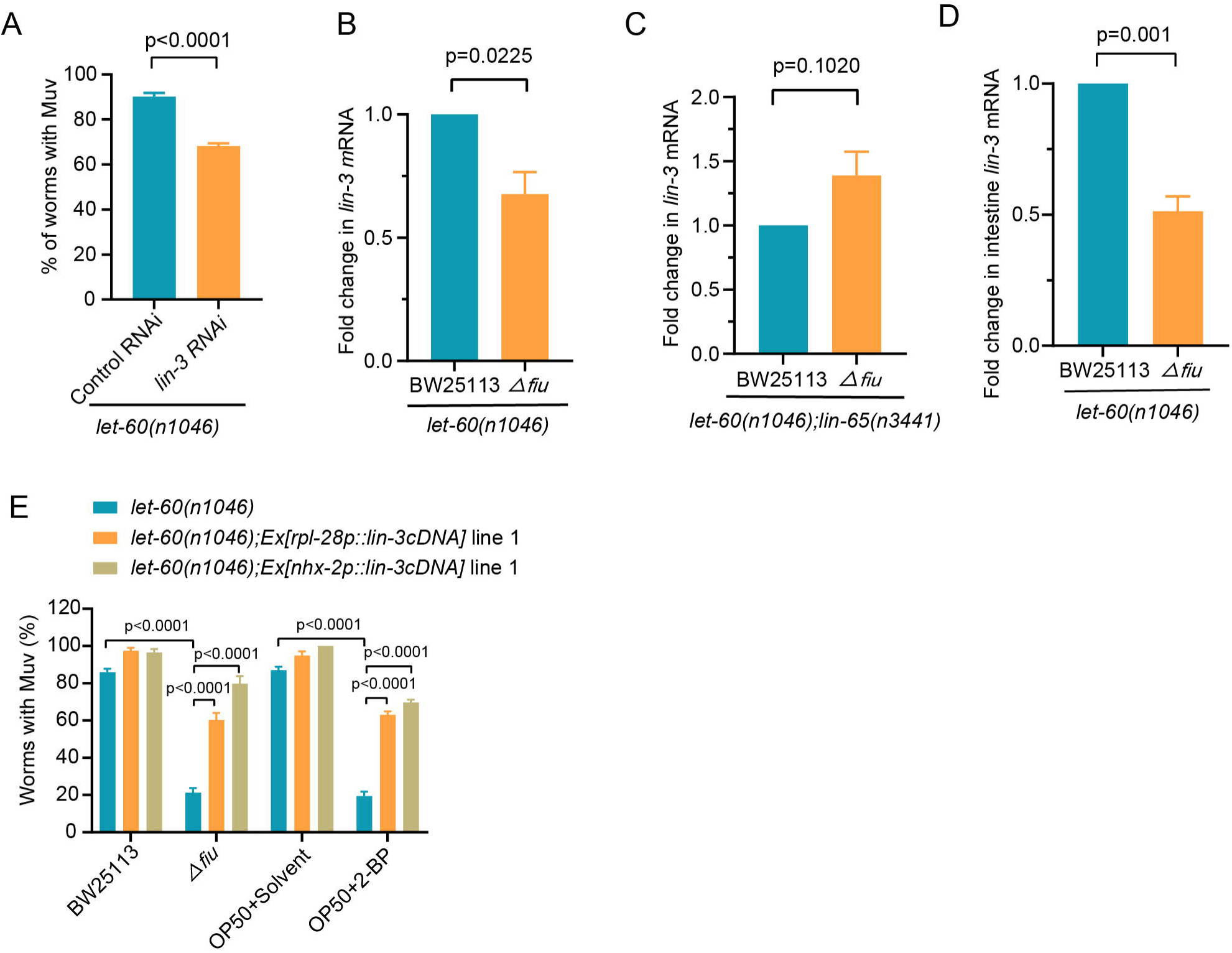
The decrease in *lin-3* transcription partially mediates LIN-65’s role in mediating the impact of Δ*fiu E. coli* on mitigating the *let-60(n1046)*-induced Muv phenotype. **(A)** A bar graph showing that *lin-3(RNAi)* partially inhibits the *let-60(n1046)*-induced Muv phenotype compared to the RNAi control group. The Muv rate is analyzed in worms subjected to the indicated treatments at the day-1 adult stage. The data are presented as the mean ± SEM by unpaired t test. **(B)** Bar graphs illustrating the decrease in *lin-3* mRNA in *let-60(n1046)* worms treated with Δ*fiu E. coli* compared to the BW25113 control group. qPCR analyses of *lin-3* mRNA were conducted at the L2-L3 stage in *let-60(n1046)* worms subjected to the specified treatments. The data are presented as the mean ± SEM by unpaired t test. **(C)** Bar graphs showing that Δ*fiu E. coli* treatment does not repress *lin-3* transcription in the absence of *lin-65*. qPCR analyses of *lin-3* mRNA were performed at the L2-L3 stage in *let-60(n1046);lin-65(n3441)* worms subjected to the specified treatments. The data are presented as the mean ± SEM by unpaired t test. **(D)** Bar graphs illustrating the significant decrease in *lin-3* mRNA in the intestines of *let-60(n1046)* worms treated with Δ*fiu E. coli*, compared to the BW25113 control. mRNA from the dissected intestines of worms with specified genotypes and treatments was isolated and analyzed using qPCR. The data are presented as the mean ± SEM by unpaired t test. **(E)** Bar graphs indicating that the suppressive effects of Δ*fiu E. coli* and 2-BP treatment on Muv are reversed by *lin-3* overexpression. The Muv rate in *let-60(n1046)* with the indicated extra-chromosomal array transgenes and treatments was assessed at the day-1 adult stage. The *rpl-28p* and *nhx-2p* promoters were utilized to overexpress *lin-3* in ubiquitous tissues and the intestine, respectively. The data are presented as the mean ± SEM by unpaired t test.

Then, we tested the potential downregulation of *lin-3* by LIN-65 in response to Δ*fiu* bacteria. Given that LIN-65 is involved in chromatin remodeling and gene silencing [38], we hypothesized that LIN-65 activation in Δ*fiu* bacteria-treated worms may suppress *Ras(gf)* effects by downregulating *lin-3* transcription. To test this hypothesis, we measured *lin-3* mRNA levels by qPCR in *let-60(gf)* worms treated with Δ*fiu E. coli* or the control BW25113 at the L2-L3 stage, during which Ras signaling is activated to specify the vulval fate of VPCs [26]. As expected, treatment with Δ*fiu* bacteria led to a decrease in *lin-3* mRNA levels in *let-60(n1046)* worms (Figure 5B). Moreover, *lin-65* is required for Δ*fiu*-induced repression of *lin-3* transcription, as evidenced by the failure of Δ*fiu* bacteria to repress *lin-3* mRNA levels in *let-60(n1046);lin-65(n3441)* worms (Figure 5C). Taken together, these results indicate that the repression of *lin-3* transcription by Δ*fiu* bacteria is dependent on *lin-65*.

Our qPCR analyses of *lin-3* transcription from the entire worm indicate approximately a 30% reduction (Figure 5B), prompting us to explore whether repression of *lin-3* transcription by Δ*fiu* bacteria treatment primarily occurs in certain tissues. Given that LIN-65 levels predominantly increased in intestinal nuclei, we investigated whether *lin-3* transcription in the intestine is significantly decreased. Using a previously established transgene, *Is[lin-3p::GFP]* [42], to analyze *lin-3* expression in *let-60(n1046)* worms at the L2-L3 stage, we found that the GFP signal was undetectable in the gut, possibly due to its diffusion in the cytoplasm of large intestinal cells. However, the GFP signal was primarily observed in anchor cells and did not decrease following Δ*fiu E. coli* treatment (Figures S5A and S5B). To quantify *lin-3* mRNA in the intestine, we dissected *let-60(n1046)* worms treated with Δ*fiu* or control BW25113 *E. coli* at the L2-L3 stage to isolate the intestine for qPCR analyses. Notably, *lin-3* transcription in the intestine was reduced by 50% (Figure 5D), in contrast to the unchanged levels in anchor cells.

To further support the critical role of intestinal *lin-3* repression in mediating the effects of Δ*fiu E. coli* and iron reduction on suppressing *let-60(n1046)*-induced Muv, we analyzed the potential of intestine-specific overexpression of *lin-3* in reversing the function of Δ*fiu E. coli* in repressing *let-60(gf)*-caused Muv. We observed that *nhx-2p* promoter-driven intestinal *lin-3* overexpression significantly reversed the suppressive effects of Δ*fiu E. coli* and 2-BP treatments on Muv in *let-60(n1046)* worms, as evidenced by the Muv rate increasing from approximately 20% to over 70% (Figure 5E and S5C). Additionally, *lin-3* overexpression using the ubiquitous promoter *rpl-28p* effectively restored the Muv phenotype in *let-60(n1046)* worms treated with either the Δ*fiu* strain or the 2-BP chelator, as indicated by an increase in the percentage of worms with the Muv phenotype from approximately 20% to 60% upon *lin-3* overexpression (Figure 5E and S5C). This similar reversing effect observed with both intestinal and ubiquitous overexpression of *lin-3* further confirms that LIN-65-mediated repression of *lin-3* transcription in the intestine is a significant contributor to the suppressive effects of Δ*fiu* bacteria on *let-60(n1046)*-induced Muv. In summary, treatment with Δ*fiu* bacteria reduces host iron availability in the mitochondria, leading to increased LIN-65 in the nuclei, which subsequently decreases *lin-3* transcription and represses *let-60(n1046)*-induced Muv.

## Discussion

Investigating whether microbes can alleviate developmental defects caused by genetic mutations and how bacterial activity can be modulated to enhance host health are important research endeavors. We conducted a comprehensive analysis and identified 151 *E. coli* genes that, when inactivated, can reverse the developmental defect, Muv, caused by a hyperactivated *Ras* mutation in *C. elegans*. These bacterial genes are involved in a variety of biological processes, indicating that the modulation of various microbial processes can inhibit host Ras signaling. Strikingly, we found that inactivation of bacterial iron-siderophore transport genes suppressed hyperactivated Ras signaling by repressing LIN-3/EGF transcription. Mechanistically, bacteria reduce the labile ferrous iron levels in the host, leading to *lin-3* transcription downregulation through an iron-perceiving mechanism involving the translocation LIN-65 to the nucleus (Figure 6). In summary, our findings describe what changes in bacterial activity can suppress *Ras(gf)* disorders and provide mechanistic insights into the modulation of Ras signaling by microbes, supporting the application of microbial activity modulation to improve host genetic defects.

**Figure 6.**
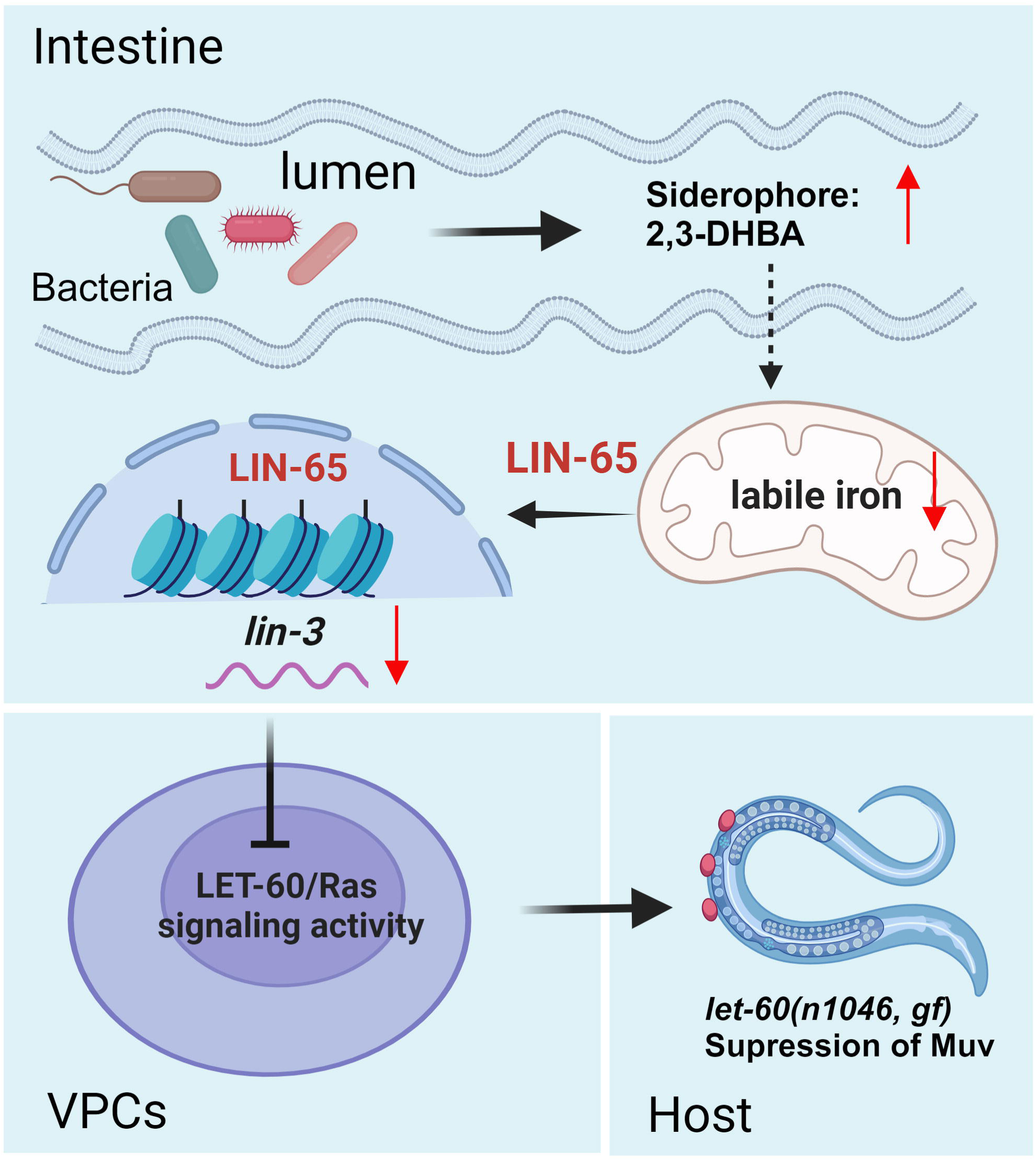
Model for the mechanism of Δ*fiu E. coli* in suppressing Muv phenotypes of *let-60(n1046)* worms. *E. coli* mutants lacking specific genes involved in iron acquisition, such as *fiu*, effectively mitigate the Muv phenotype in *let-60(n1046) C. elegans*, which carries a gain-of-function mutation in Ras. These Δ*fiu E. coli* secrete higher levels of the 2,3-DHBA siderophore compared to the wild-type strain, leading to a reduction in host mitochondrial iron levels. In response to the decreased iron availability induced by the bacteria, the host chromatin modifier LIN-65 translocates to the nucleus to suppress *lin-3/EGF* transcription, thereby repressing hyperactive let-60/*Ras(gf)*-induced Muv development.

How cells perceive the level of essential iron and then orchestrate the growth signaling state to adapt to iron availability is not examined. Our results revealed a mechanism of iron-perceiving, which transduces iron levels into Ras signaling through EGF modulation and involves both mitochondria and the Lin-65. Our study thus elucidates a novel function of the SynMuv gene *lin-65* in mediating the role of iron in regulating Ras signaling. This newly discovered mechanism likely ensures that Ras signaling is downregulated to inhibit cell proliferation in response to low iron levels. Interestingly, Ras signaling can also modulate iron levels by regulating iron metabolism genes [43], thus indicating that reciprocal regulation between iron and Ras signaling occurs. These mechanisms may enable precise coordination between iron availability and growth-related signaling pathways.

Our discovery that gut microbes can suppress the overactivation of *Ras* signaling in the host organism by decreasing iron availability offers novel perspectives on the prevention of *Ras(gf)*-induced disorders, such as tumors. A few studies have suggested that high iron levels are significantly associated with the occurrence and progression of cancer [44], such as colorectal cancer and pancreatic cancer. These observations suggest that sufficient iron is important for *Ras(gf)*-mediated activation of downstream signaling, suggesting that decreasing iron levels could inhibit tumorigenesis. Indeed, clinical studies have shown that certain cancer patients respond to iron chelating agent treatment [44], but the mechanism through which the iron reduction suppresses tumor growth remains relatively unexplored. Therefore, our study on iron insufficiency-mediated suppression of Ras signaling may provide a mechanistic understanding of how tumor growth can be inhibited by reducing iron levels, supporting the potential of developing iron-targeting therapies.

The identification of microbe mutants that suppress hyperactivated Ras signaling opens up potential avenues for treating *Ras(gf)*-associated defects through modulating bacterial activities. The microbiota have been suggested to play critical roles in tumor initiation and development, with certain microbes exerting anticancer effects [45]; however, the precise molecular pathways involved are poorly understood. Our study provides mechanistic understanding of the anti-Ras oncogenic signaling effect of the microbiota, which sheds light on the development of microbial-based approaches for cancer therapy. For example, targeting bacterial iron acquisition systems with antibiotics may hold promise for suppressing *Ras(gf)*-induced tumors. Moreover, our genome-wide screen revealed 151 *E. coli* gene mutations that can suppress hyperactivated Ras signaling, thus providing a valuable resource for further investigation of the mechanisms underlying microbe-mediated suppression of oncogenic Ras. In summary, our findings demonstrate that microbes can suppress the effects of host genetic defects, which expands our understanding of the role of microbes in promoting host health and offers valuable insights for the development of effective approaches to manage *Ras(gf)*-induced defects using microbial interventions.

## Acknowledgments

We thank the CGC (funded by NIH [P40OD010440]) for strains, Ye Tian’s laboratory (IGDB, CAS) for strains, and all the members of the Tang lab for comments and suggestions. This work was supported by the National Natural Science Foundation of China (No. 32350015, No. 32070565, and No. 31871465), the National Key Research and Development Program of China (No. 2019YFA0802900), the HRHI program (No. 202209003 and No. 202109007) of the Westlake Laboratory of Life Sciences and Biomedicine, the Zhejiang Provincial Natural Science Foundation of China (XHD24C0701 and No. LQ23C040002), the Westlake Education Foundation of Westlake University and the Zhejiang Provincial Key Laboratory Construction Project.

## Author contributions

H.T. conceptualized and supervised the study. M.D. conceived, designed, and performed most of the experiments as well as analyzed the data. Y.W. performed part of the iron determination assays and analyses. G.L. provided technical support for the iron determination with staining. M.D. and H.T. wrote the manuscript.

## Declaration of interests

The authors declare no competing interests.

## Supplemental figure legends

**Figure S1.**
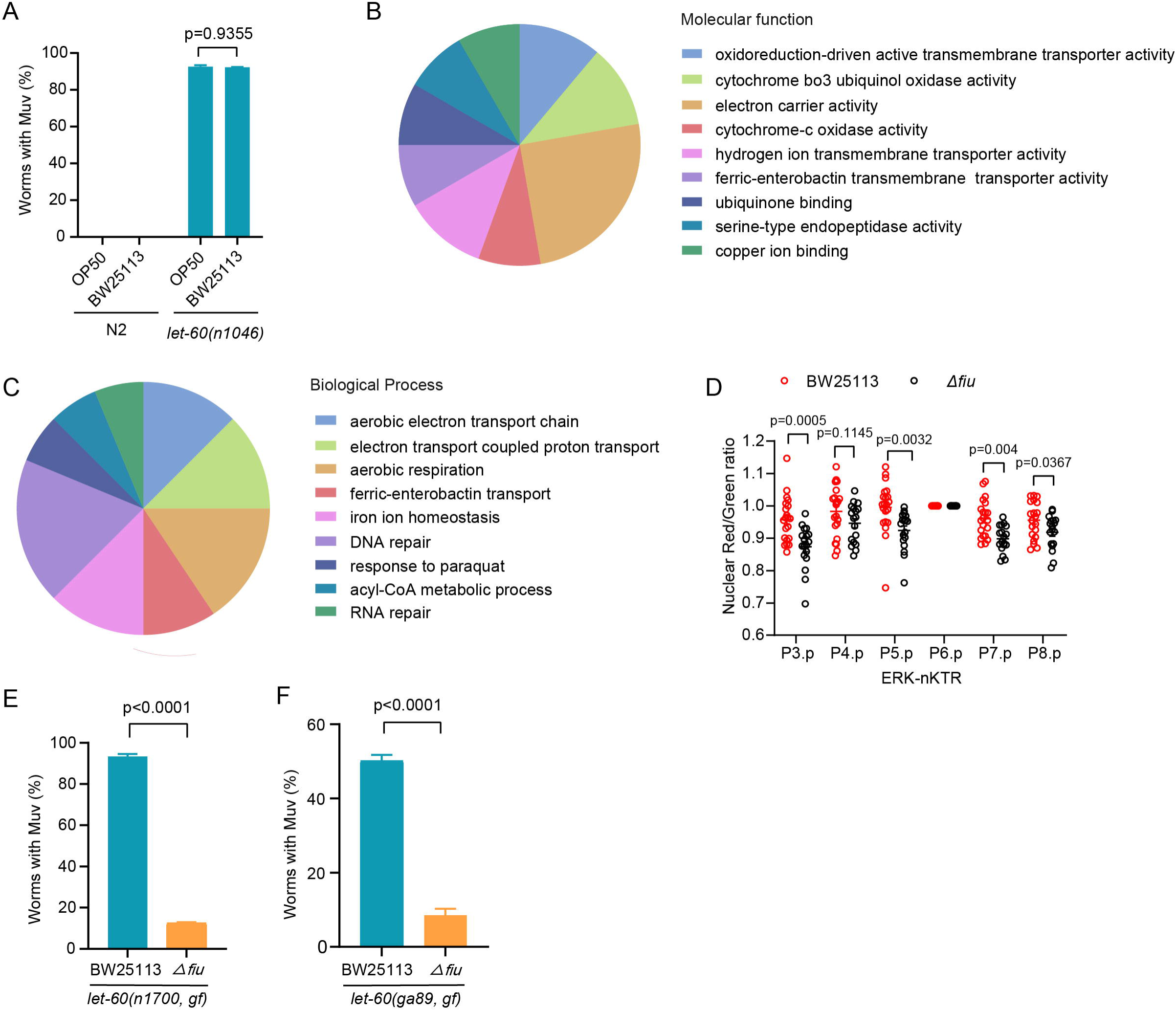
Inactivation of various bacterial processes suppresses *let-60(gf)*-induced Muv and Δ*fiu E. coli* treatment represses Ras signaling. **(A)** Bar graph showing the percentage of Muv in N2 and *let-60 (n1046)* worms with indicated bacterial treatment. The L1-staged worms grown on either OP50 or BW25113 were scored for Muv when reaching day-1 adulthood. The data are presented as the mean ± SEM. ns, p = 0.9355 as determined by two-way ANOVA. **(B-C)** Gene Ontology analyses. Analyses of biological processes and molecular function of the 151 *E. coli* genes identified in the screen in Figure 1B revealed that these genes widely participate in different processes, including bacterial iron transportation. **(D)** Graphs indicating that MPK-1/ERK activity is decreased in the VPCs. MPK-1 activity is assessed using a previously established kinase translocation reporter that expresses *arTi85[lin-31p::ERK-KTR-mClover::T2A::mCherry-H2B]*, by calculating the nuclear Red/Green Ratio. The nuclear Red/Green Ratio was scored at the L2-L3 stage in *let-60(n1046)* treated with the indicated bacteria. Each dot represents the ratio of Red/Green fluorescence intensity. n=21 for BW25113 treatment and n=18 for Δ*fiu E. coli* treatment. **(E-F)** Bar graphs illustrate that Δ*fiu E. coli* treatment suppresses the Muv phenotypes induced by two additional *let-60(gf)* alleles, *let-60(n1700)* and *let-60(ga89)*. The Muv rate of *let-60(n1700)* and *let-60(ga89)* worms treated with BW25113 and Δ*fiu E. coli* was scored. Data are presented as the mean ± SEM. p < 0.0001 by unpaired t-test.

**Figure S2.**
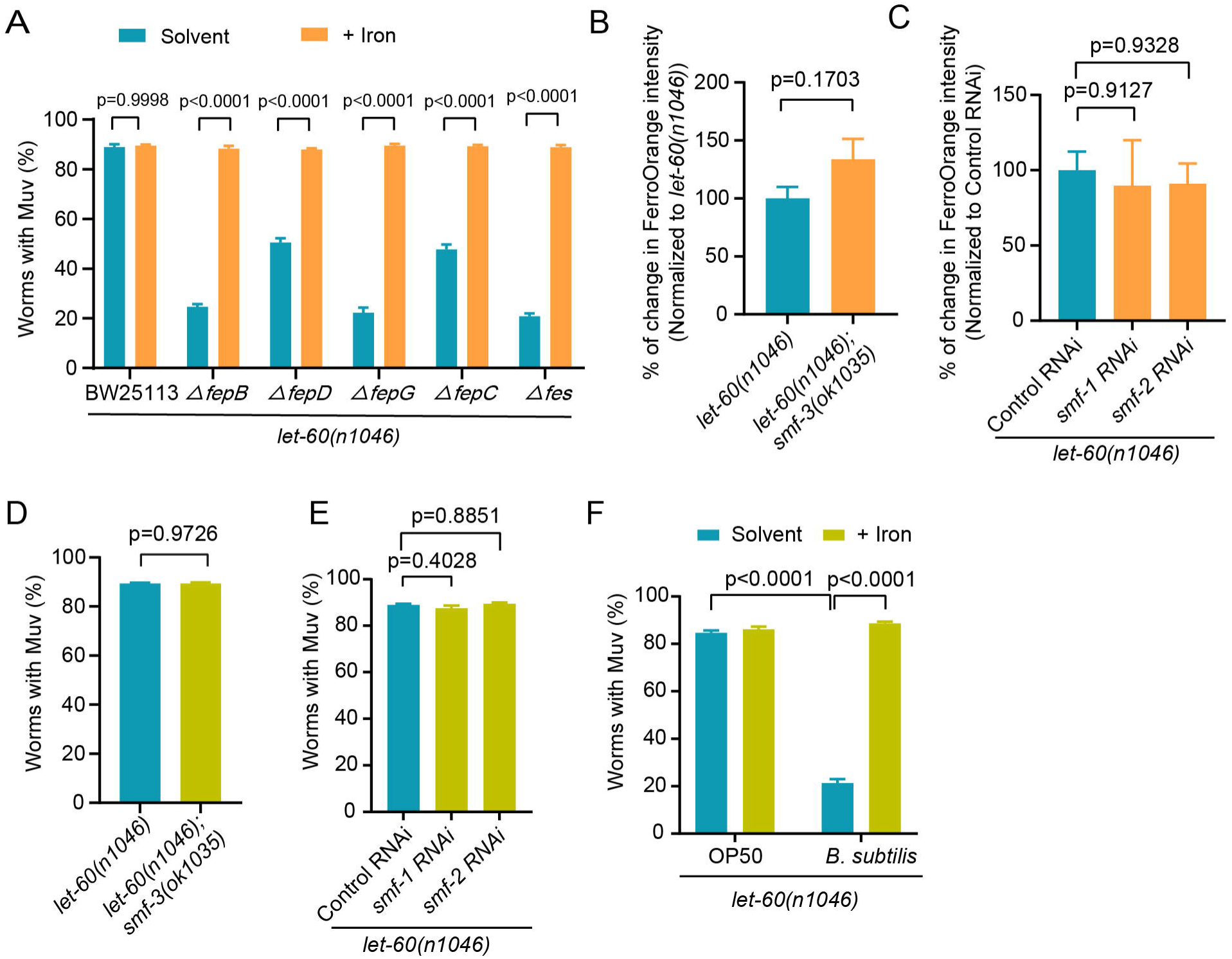
Bacteria, including *E. coli* with mutation in iron acquisition and *B. subtilis*, that decrease host iron availability, can suppress *let-60(gf)*-induced Muv. **(A)** Bar graphs demonstrate that iron supplementation reverses the suppressive effect of bacteria with mutations in genes involved in Fe-siderophore transport on the *let-60(n1046)*-induced Muv phenotype. The Muv rate is analyzed in *let-60(n1046)* worms treated with the specified *E. coli* mutants, with or without FeCl_3_ supplementation, at the day-1 adult stage. The data are presented as the mean ± SEM determined by two-way ANOVA. **(B)** Bar graphs illustrate that the levels of ferrous iron, as stained with FerroOrange, remain unchanged between *let-60(n1046)* and *let-60(n1046);smf-3(ok1035)* worms. FerroOrange intensity was analyzed in the specified worms grown on OP50 at the day-1 adult stage. The data are presented as the mean ± SEM according to the unpaired t test. **(C)** Bar graphs demonstrate that the levels of ferrous iron, stained with FerroOrange, remain unchanged in worms treated with *smf-1(RNAi)* or *smf-2(RNAi)*. The fluorescence intensity of FerroOrange was analyzed in *let-60(n1046)* worms treated with the specified RNAi at the day-1 adult stage. The data are presented as the mean ± SEM by one-way ANOVA. **(D)** Bar graphs showing that the Muv rate remains unchanged in the *let-60(n1046)* and *let-60(n1046);smf-3(ok1035)* worms. The Muv phenotype was scored in the indicated worms grown on OP50 at day-1 adult stage. The data are presented as the mean ± SEM, determined by unpaired t test. **(E)** Bar graphs indicating that the Muv rate in the *let-60(n1046)* worms were not changed after *smf-1(RNAi)* or *smf-2(RNAi)* treatment. The Muv rate was analyzed in *let-60(n1046)* worms treated with the specified RNAi at the day-1 adult stage. The data are presented as the mean ± SEM by one-way ANOVA. **(F)** Bar graphs illustrate that *B. subtilis* treatment suppresses *let-60(gf)*-induced Muv phenotype, which can be reversed by iron supplementation. The Muv rate was analyzed in *let-60(n1046)* worms treated with *B. subtilis* or OP50, with or without FeCl_3_ supplementation. The data are presented as the mean ± SEM determined by two-way ANOVA.

**Figure S3.**
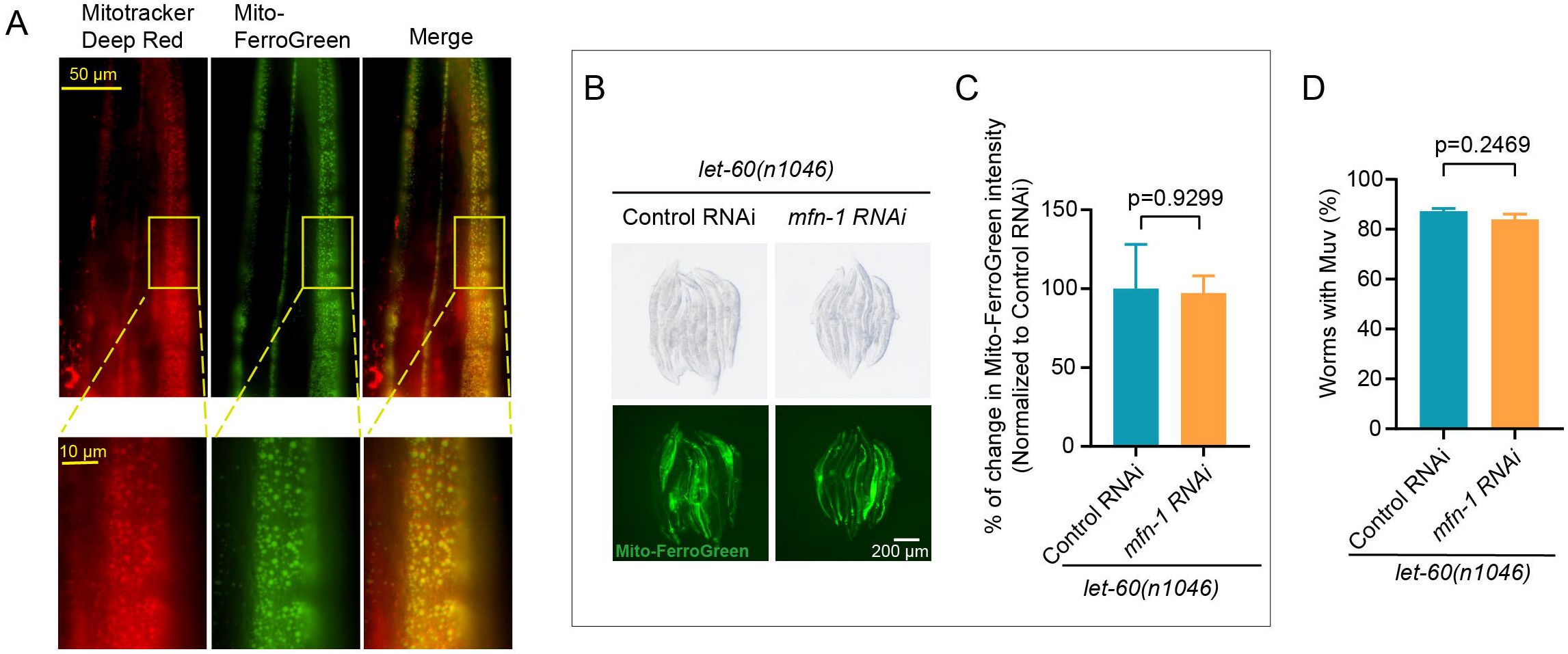
Mito-FerroGreen stains mitochondrial ferrous iron and the *mfn-1(RNAi)* is not sufficient to decrease mitochondrial iron and suppress Muv. **(A)** Microscopic images showing the mitochondrial localization of Mito-FerroGreen staining, as indicated by its colocalization with MitoTracker Deep Red. **(B-C)** Images and bar graphs showing that *mfn-1(RNAi)* alone is not sufficient to decrease the mitochondrial ferrous iron level. Intensity of Mito-FerroGreen in *let-60(n1046)* worms with indicated RNAi treatment was analyzed. The data are presented as the mean ± SEM by unpaired t test. **(D)** Bar graphs indicating the Muv rate of *let-60(n1046)* worms remained unchanged after *mfn-1* RNAi treatment. The Muv rate is analyzed at day-1 adult stage. The data are presented as the mean ± SEM by unpaired t test.

**Figure S4.**
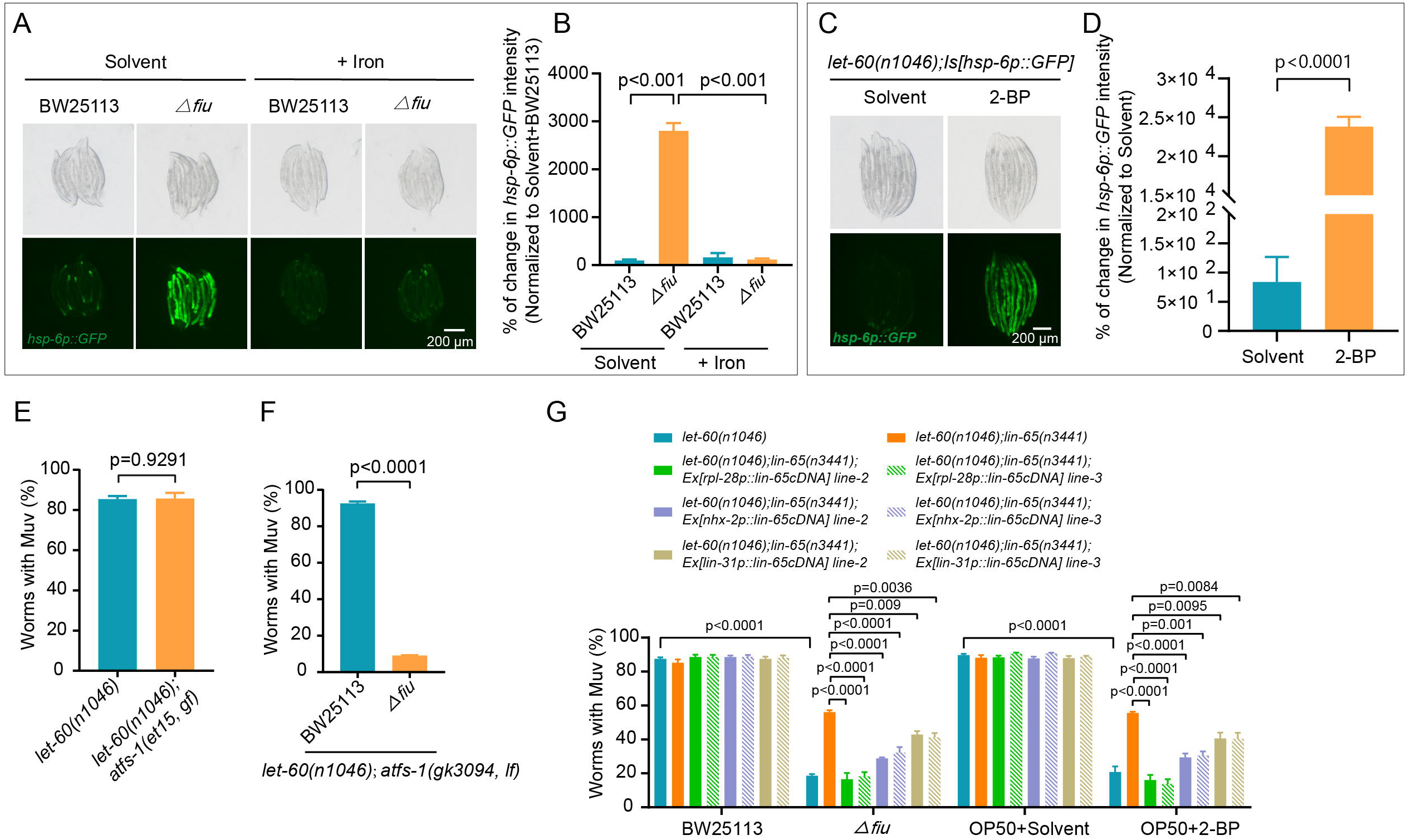
Δ*fiu* bacteria-induced reduction of iron availability triggers the mitochondrial unfolded protein response (mito^UPR^), and the mito^UPR^ regulator LIN-65 mediates the role of Δ*fiu* bacteria in suppressing *let-60(lf)*-caused Muv. **(A-B)** Images and bar graphs illustrate the induction of the mitochondrial stress reporter, *hsp-6p::GFP*, and its reversal by iron (FeCl_3_) supplementation. The GFP signal was analyzed in *let-60(n1046);Is[hsp-6p::GFP]* worms subjected to specified treatments. The data are presented as the mean ± SEM by one-way ANOVA. **(C-D)** Images and bar graphs illustrate the induction of the mitochondrial stress reporter, *hsp-6p::GFP*, by iron chelator 2-BP treatment. The GFP signal in *let-60(n1046);hsp-6p::GFP* worms treated with 2-BP or solvent was analyzed. The data are presented as the mean ± SEM by unpaired t test. **(E)** Bar graphs illustrate that the Muv rate remained unchanged in *let-60(n1046)* worms upon introducing the *atfs-1(et115, gf)* allele. The Muv rate was analyzed in worms with the indicated genotypes grown on OP50 at the adult day-1 stage. The data are presented as the mean ± SEM by unpaired t test. **(F)** Bar graphs showing that the suppressive effect of Δ*fiu* bacteria on *let-60(n1046)-*induced Muv is independent of *atfs-1*. The Muv rate in *let-60(n1046);atfs-1(gk3094)* worms, following specified bacterial treatments, was analyzed. The data are presented as the mean ± SEM by unpaired t test. **(G)** Bar graphs indicating the Muv rate of *let-60(n1046);lin-65(n3441)* worms carrying the specified transgenes and treated with designated bacteria and 2-BP. Consistent with the results in Figure 4B, two other independent extra-chromosomal array transgenic lines expressing *lin-65* driven by the promoters *rpl-28p*, *nhx-2p*, and *lin-31p* were able to partially recover the suppressive effect of Δ*fiu E. coli* and 2-BP on *let-60(n1046)-*induced Muv in *lin-65(lf)* mutant worms. The data are presented as the mean ± SEM by one-way ANOVA.

**Figure S5.**
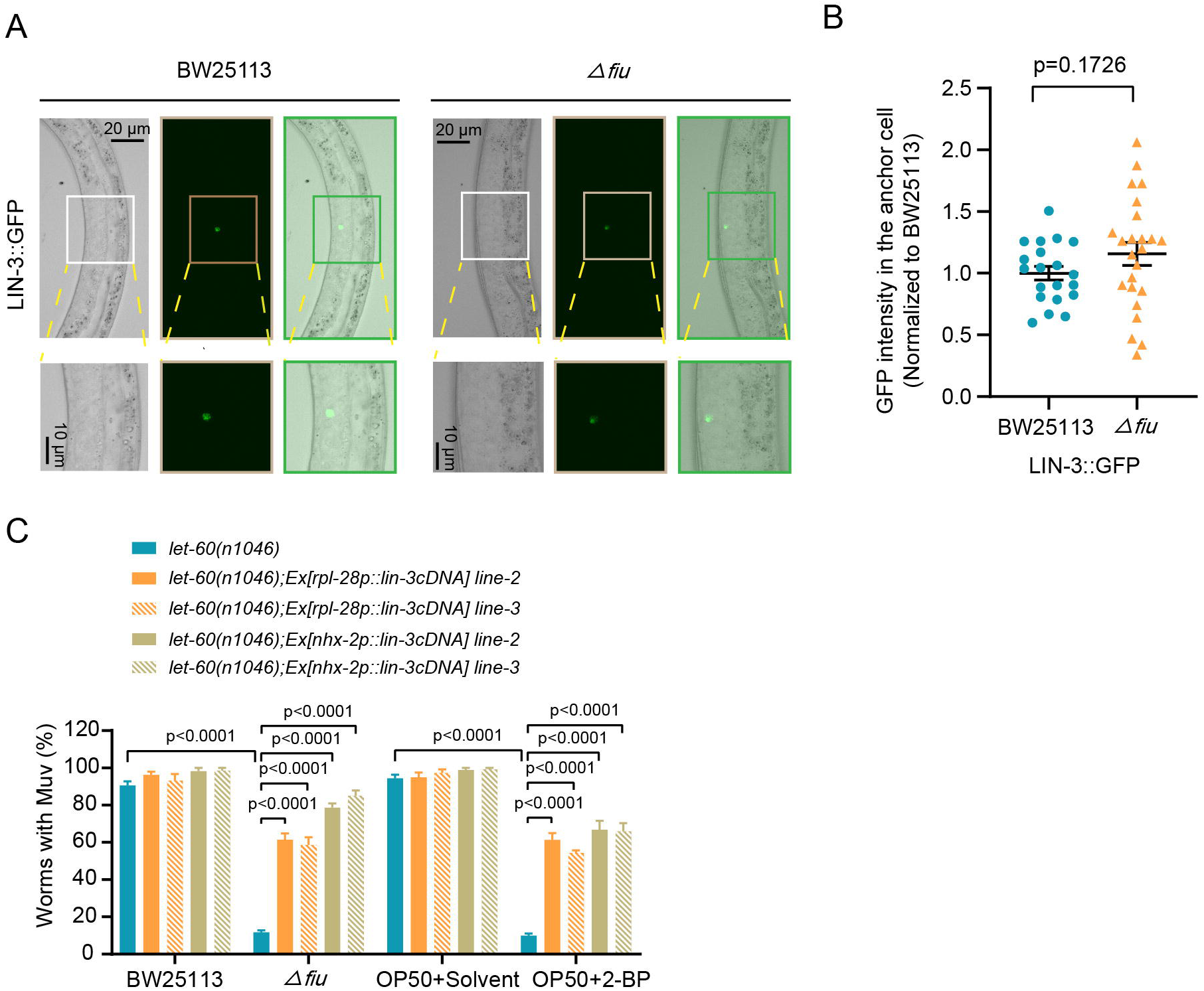
Intestine-specific over-expression of *lin-3* reverses the impact of Δ*fiu* bacteria and iron insufficiency on repressing *let-60(gf)*-induced Muv. **(A-B)** Microscopic images and bar graphs demonstrate that Δ*fiu E. coli* treatment does not decrease LIN-3 levels in the anchor cell. At the L2-L3 stage, the LIN-3::GFP signal intensity in the anchor cell was analyzed in the *let-60(n1046);syIs107 [lin-3(delta pes-10)::GFP + unc-119(+)]* worms treated with the specified bacteria. The data are presented as the mean ± SEM by unpaired t test. **(C)** Bar graphs depict the Muv rate of *let-60(n1046)* worms overexpressing *lin-3* in various tissues following treatment with designated bacteria and 2-BP. Consistent with the findings in Figure 5E, two additional independent extrachromosomal array transgenic lines, expressing *lin-3* under the control of the *rpl-28p* and *nhx-2p* promoters, partially reversed the suppressive effects of Δ*fiu E. coli* and 2-BP on *let-60(n1046)*-induced Muv. The data are presented as the mean ± SEM by unpaired t test.

## STAR Methods

### Key Resources Table

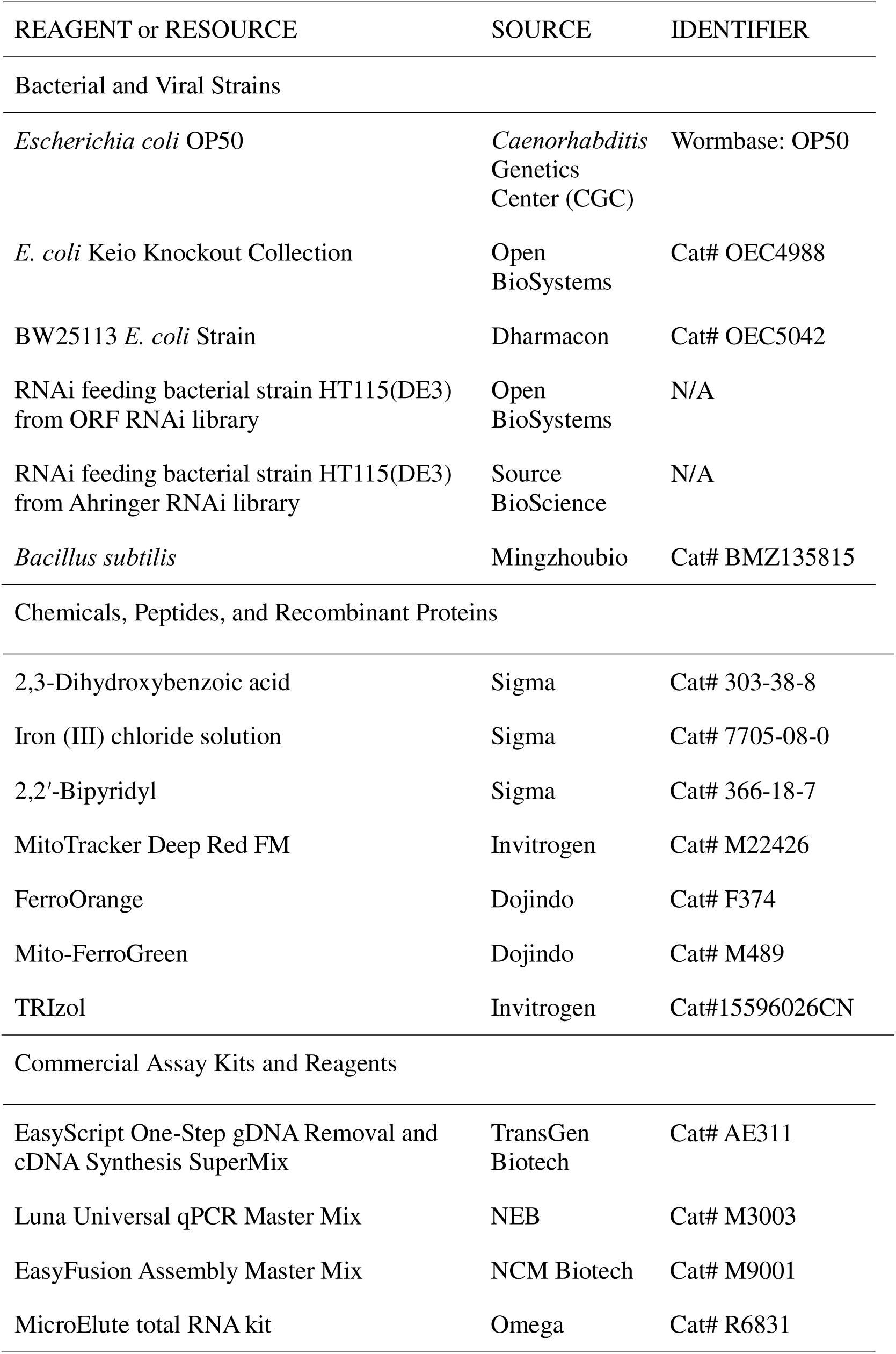
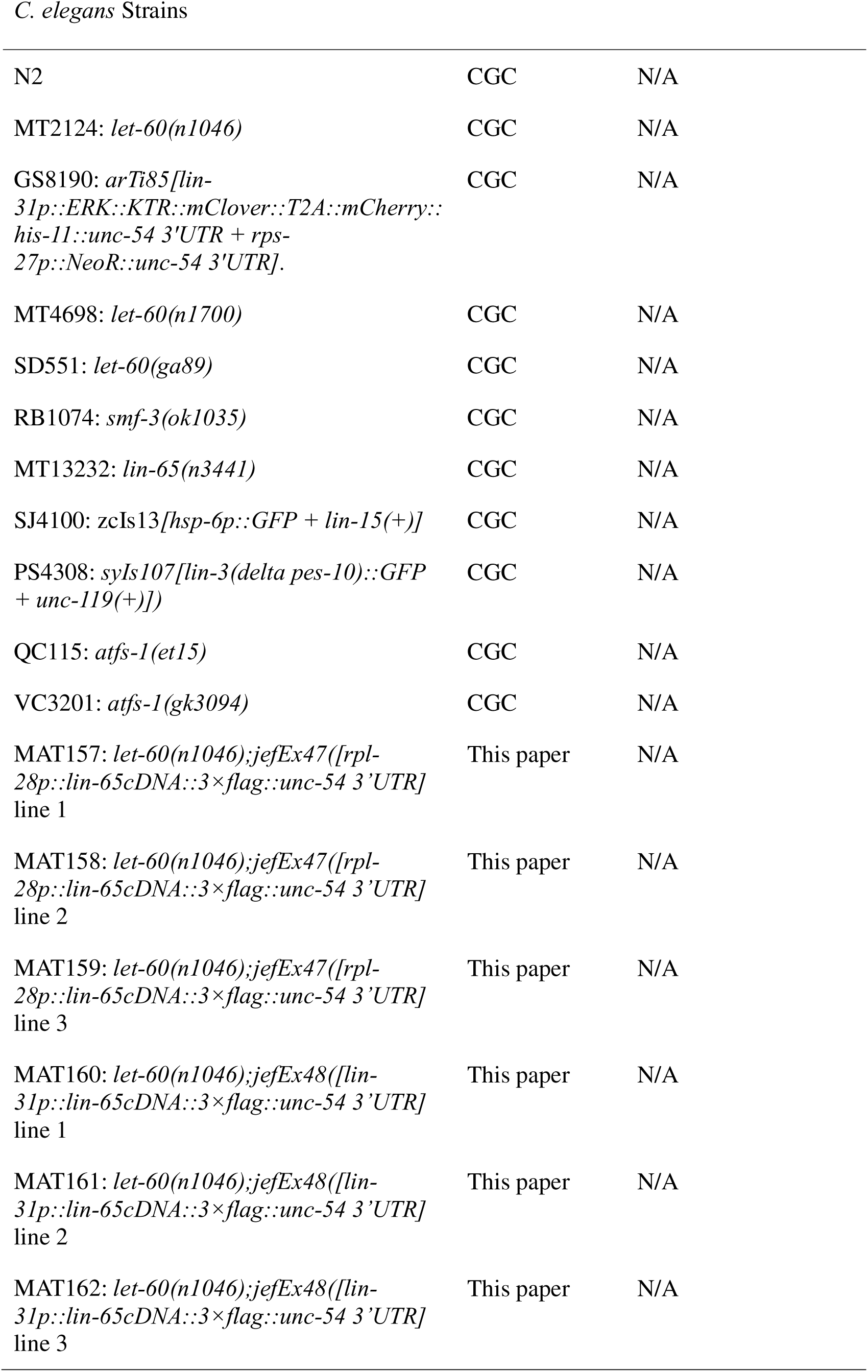
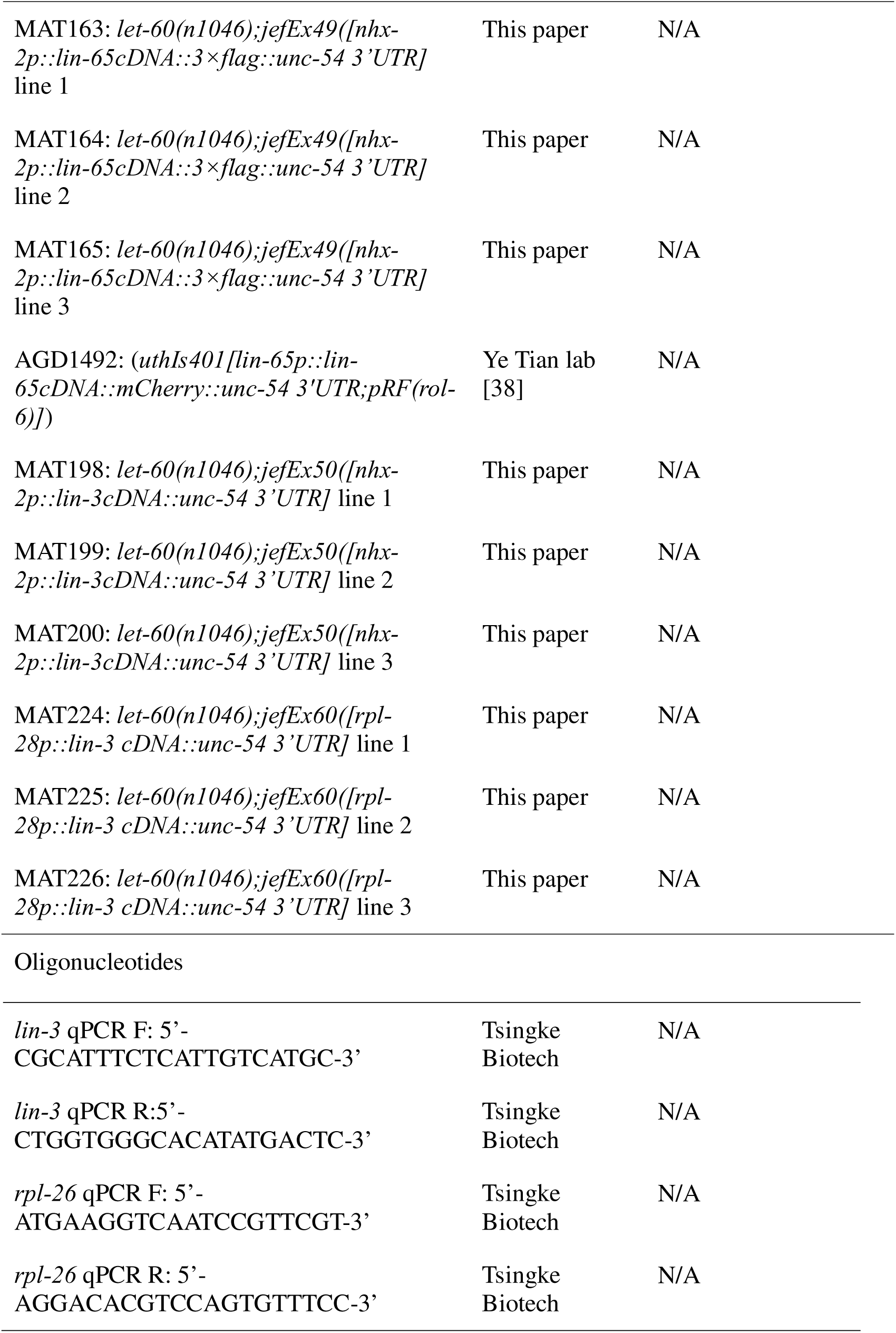

### *C. elegans* strains and maintenance

*C. elegans* strains, listed in the key resource table, were maintained on nematode growth medium (NGM) plates with OP50 bacteria at 20°C. The *uthIs401* strain was provided by the Ye Tian laboratory [38]. All the other strains were obtained from the *Caenorhabditis* Genetics Center (CGC).

### Microbial strains and maintenance

The *E. coli* Keio mutant strains were cultured overnight at 37°C in Luria–Bertani (LB) medium supplemented with 25 μg/mL kanamycin. Subsequently, they were spread on NGM plates supplemented with 25 μg/mL kanamycin for further use. *B. subtilis* bacteria were cultured overnight at 30°C in LB medium and spread on NGM plates for further use.

### Screening the *E. coli* Keio knockout collection for Muv suppression in *C. elegans*

Synchronized L1 *let-60(n1046)* worms were added to the prepared *E. coli* Keio mutant bacteria plates and cultured at 20°C. The percentage of worms with the Muv phenotype, displaying as extra vulva, was scored on day 1 adulthood. The entire knockout library containing 3985 *E. coli* mutation was used in the primary screen to identify potential bacterial mutants that can repress the Muv phenotype of in *let-60(n1046)* worms, and 350 positive hits were obtained. Then, these 350 candidates were screened three more times to further confirm their ability to suppress Muv.

### Scoring Vulval induction

VPCs induction, characterized by invagination, were scored at the L4 stage through using the transgene *lin-31p:::mCherry::H2B* that highlights the nucleus of the VPCs. In wild-type worms, P5.p, P6.p, and P7.p VPCs adopt vulva fate, thus counting as 3 VPCs with induction [26]. If the P3.p, P4.p, and P8.p VPCs overdifferentiate and undergo invagination, their cell type changes, and each differentiation is counted as 1 induced VPC. If P5.p and P7.p undergo overdifferentiation and invagination, each cell of this differentiation is considered a 0.5-fold increase in the VPC [46].

### Analysis of MPK-1/ERK activity with a kinase translocation reporter

The ERK activity in the VPCs was analyzed with a previously-established kinase translocation reporter, *lin-31p::ERK-KTR-mClover::T2A::mCherry-H2B*, which converts phosphorylation state into a nucleocytoplasmic localization within the cell [17]. Specifically, L2-L3 stage larvae with indicated genotypes and treatments were mounted on the 2% agarose pad, Z-stacks of mClover and mCherry fluorescence were captured simultaneously using the confocal microscope (Olympus). The same photograph settings were adopted for all the images, the images were analyzed with the Imaris, in which under 3D view conditions, surfaces were created for the image and mCherry signal was chosen for the source channel to render the image, the mean intensity of the mCherry (Red) and mClover (Green) signal for the six VPCs were recorded to calculate the nuclear Red/Green Ratio that represents the MPK-1/ERK activity.

### Quantification of 2,3-DHBA generated by the bacteria

Bacteria secret siderophores to the extracellular space for acquiring iron, thus 2,3-DHBA siderophore in the culture medium were analyzed. Specifically, BW25113 and Δfiu bacterial clones were inoculated and cultured in LB medium over night at 37°C. After centrifugation of 1 mL bacterial culture, 100 μL of the supernatant was collected and then 900 μL of methanol was added, and the mixture was kept at −80°C for 24 hours to allow the extraction of 2,3-DHBA. Subsequently, the samples were dried at 4°C with a vacuum centrifugal concentrator (Christ, China) and dissolved in 50 μL of methanol. 2,3-DHBA was quantified via mass spectrometry (Agilent, USA), and commercially obtained 2,3-DHBA was used as the standard sample. The amount of 2,3-DHBA was normalized to the bacterial content, which was determined by counting the number of clones produced by 2 μL of the bacterial culture medium.

### Chemical supplementation

The 2,3-dihydroxybenzoic acid (2,3-DHBA), 2,2′-bipyridyl (2-BP), and FeCl_3_ were supplemented to manipulate the iron levels in the worms. These chemicals were dissolved in water or DMSO to generate the stock solutions, which were added to the NGM plates with the final concentration at 18 mg/mL for 2,3-DHBA, 50 μM for 2-BP, and 4 mM for FeCl_3_.

### Iron determination in worms

FerroOrange and Mito–FerroGreen were used, respectively, to detect total Fe^2+^ in worms and total Fe^2+^ in the mitochondria of the worms. These Fe^2+^ indicators were freshly prepared according to the manufacturer’s instructions. About 30 day-1 adult worms with indicated treatments were stained with FerroOrange (10 μM) and Mito–FerroGreen (100 μM) for 2 hours in M9 buffer. For the Mito-Tracker DeepRed and Mito-FerroGreen co-staining, the worms were stained with Mito-FerroGreen as previously described, washed with M9 buffer, dissected, and subsequently stained with Mito-Tracker Deep Red for 30 minutes. These worms were randomly collected for fluorescence imaging.

### Transgene construction

These transgenes were generated in this study:

*let-60(n1046);lin-65(n3441);Ex[rpl-28p::lin-65cDNA::3×Flag::unc-54 3’UTR]*

*let-60(n1046);lin-65(n3441);Ex[nhx-2p::lin-65cDNA::3×Flag::unc-54 3’UTR]*

*let-60(n1046);lin-65(n3441);Ex[lin-31p::lin-65cDNA::3×Flag::unc-54 3’UTR]*

*let-60(n1046);Ex[rpl-28p::lin-3cDNA::unc-54 3’UTR]*

*let-60(n1046);Ex[nhx-2p::lin-3cDNA::unc-54 3’UTR]*

To achieve tissue-specific expression of *lin-65* in *let-60(n1046); lin-65(n3441)* worms, the *lin-65* cDNA (2184 bp), fused with a 3×Flag tag at the C-terminus, was amplified via PCR using primers that correspond to the *lin-65* regions and incorporate the 3×Flag sequence. The obtained sequence was inserted into pBluescript SK (pBSK), to produce the *lin-65 cDNA ::3×Flag::unc-54 3’UTR* plasmid. Then, the *rpl-28* promoter (1437 bp), *nhx-2* promoter (1971 bp), and *lin-31* promoter (4544 bp) sequences were amplified via PCR and subcloned into the upstream of *lin-65 cDNA::3×Flag::unc-54 3’UTR,* to create the *rpl-28p::lin-65 cDNA::3×Flag::unc-54 3’UTR*, *nhx-2p::lin-65 cDNA::3×Flag::unc-54 3’UTR*, and *lin-31p::lin-65 cDNA::3×Flag::unc-54 3’UTR* plasmids.

*lin-3* cDNA was used for tissue-specific overexpression in *let-60 (n1046)* worms*. lin-3* cDNA (1272 bp) was amplified via PCR and subcloned into the pBSK plasmid to obtain the *lin-3cDNA::unc-54 3’UTR* plasmid. Then, the *rpl-28* promoter (1437 bp) and *nhx-2* promoter (1971 bp) were inserted into the upstream of *lin-3cDNA::unc-54 3’UTR*, respectively, to obtain the *rpl-28p::lin-3cDNA::unc-54 3’UTR* and *nhx-2p::lin-3cDNA::unc-54 3’UTR* plasmids.

These obtained plasmids were used to generate extra-chromosomal array transgenic strains as previously described [47]. Specifically, the DNA mixture containing target constructs (50 ng/μL) and the co-injection marker (20 ng/μL) were microinjected into the indicated worms to generate the corresponding transgenic worms and three independent lines were obtained.

### *hsp-6p::GFP* intensity analysis

Synchronized L1-staged worms carrying *hsp-6p::GFP* were added to the plates with indicated treatment. When reaching day 1 adulthood, worms were randomly collected, and their GFP fluorescence was captured with the stereo microscope (Nikon).

### RNA isolation and real-time PCR

Worms at L2-L3 stage were harvested for analyzing the *lin-3* mRNA level. Specifically, the *let-60(n1046)* worms with indicated bacterial treatment were collected at L2-L3 stage, which, after wash with M9 buffer, were resuspended in TRIzol reagent (Invitrogen) and homogenized. Total RNA was isolated by chloroform extraction and ethanol precipitation. For extracting mRNA from the intestine, the worms with indicated treatment were dissected to release the intestine, which were collected and then used for RNA extraction by using the micro total RNA kit (Omega). The cDNA was synthesized by reverse transcription (TransGen, One-Step gDNA Removal and cDNA Synthesis SuperMix). The primers used for *lin-3* qPCR were 5’-CGCATTTCTCATTGTCATGC-3’ and 5’-CTGGTGGGCACATATGACTC-3’. Real-time PCR was performed using Universal qPCR Master Mix (NEB). The values were normalized to those of *rpl-26* (forward primer: ATGAAGGTCAATCCGTTCGT; reverse primer: AGGACACGTCCAGTGTTTCC) as the internal controls for gene expression.

### RNAi treatment

Worms were subjected to RNAi by feeding. Specifically, the bacteria for the worm gene RNAi experiments were selected from the ORF RNAi collection or the Ahringer RNAi library, which were then cultured in LB medium supplemented with 100 μg/mL ampicillin and 25 μg/mL tetracycline at 37°C overnight and spotted on NGM plates supplemented with 100 μg/mL ampicillin and 2 mM isopropyl-beta-D-1-thiogalactopyranoside (IPTG). These plates were incubated overnight at room temperature to allow IPTG-mediated induction of dsRNA expression. L1s were raised on designated RNAi bacterial plates, and when reaching day-1 adulthood, the Muv phenotype or the iron levels were scored as mentioned above.

### Quantification and statistical analysis

To reduce potential bias, worms were randomly picked under a dissection microscope for further analyzing indicated phenotypes. A minimum of three independent biological experiments were performed to ensure robustness and reliability, and the data are presented as means ± standard errors of the mean (SEMs) to illustrate the level of variation. The unpaired t-test for two groups, one-way ANOVA for more than two groups, and two-way ANOVA for more than two groups within multiple subgroups were conducted using GraphPad Prism to assess statistical significance. In all cases, p values < 0.05 were considered significant.

## Notes

### Competing Interest Statement

The authors have declared no competing interest.

